# Antimicrobial-resistance plasmids encode Evangelion, a widespread DUF4062 anti-phage defence family

**DOI:** 10.64898/2026.06.25.731849

**Authors:** Rodrigo Ibarra-Chavez, Aa Haeruman Azam, Kotaro Chihara, Amaru Miranda Djurhuus, Kristine A. Gottlieb, Azumi Tamura, Thomas Sicheritz-Pontén, Jonas S. Madsen, Kotaro Kiga

## Abstract

Many bacterial anti-phage defence systems are encoded by mobile genetic elements, yet much of this antiviral repertoire remains undiscovered. Antimicrobial-resistance plasmids are well known for disseminating antibiotic-resistance genes, but their contribution to bacterial immunity remains largely unexplored. Using phenotype-guided interrogation of a naturally occurring methicillin-resistant *Staphylococcus aureus* plasmid, we identify Evangelion, a widespread family of single-gene anti-phage defence systems in which a conserved DUF4062 core is coupled to a diversified auxiliary region required for defence activity. Evangelion systems are enriched on antimicrobial-resistance plasmids but are distributed across diverse bacterial hosts and mobile genetic elements, revealing an evolutionarily conserved defence architecture that has diversified through horizontal gene transfer and adaptation to distinct phage environments. Genetic, structural and evolutionary analyses support a model in which Eva01 senses intracellular phage replication-associated processes and couples activation of a DUF4062 effector to NAD⁺ depletion and abortive infection. Together, our findings define a previously unrecognised family of mobile anti-phage defence systems, establish DUF4062 proteins as a new component of the bacterial anti-phage repertoire, and demonstrate that phenotype-guided interrogation of mobile genetic elements provides a powerful strategy for discovering defence systems beyond the reach of current computational approaches.

## Introduction

Mobile genetic elements (MGEs) are major drivers of horizontal gene transfer (HGT) and bacterial evolution, mediating the transfer of genes and competing with one another for persistence (Frost et al., 2005; Smillie et al., 2010). Plasmids, prophages, phage satellites, integrative elements and other MGEs can carry functions that benefit their bacterial hosts while also promoting their own maintenance and spread. One important example is anti-phage immunity. Many bacterial defence systems are encoded by MGEs, allowing them to protect host cells from viral predation and to shape the distribution of competing genetic elements, including conflicts between plasmids and other MGEs (Doron et al., 2018; Makarova et al., 2011; Bondy-Denomy et al., 2016; Dedrick et al., 2017; Fillol-Salom et al., 2022; Rousset et al., 2022; Botelho et al., 2023; Pfeifer et al., 2022). However, the role of different MGE types to phage defence, and the extent to which clinically important MGEs alter phage susceptibility, remain poorly understood.

Amongst the diverse repertoire of MGEs, plasmids are especially important because they act as major reservoirs of antimicrobial resistance genes, adapt rapidly to selective pressures and spread across diverse multidrug-resistant bacteria, including the ESKAPE pathogens (Partridge et al., 2018; San Millan, 2018; Ares-Arroyo et al., 2023; GBD Antimicrobial Resistance Collaborators et al., 2024; Cazares et al., 2025). Recent studies have further highlighted plasmids as reservoirs of anti-phage and anti-defence functions, including restriction-modification systems, abortive infection systems, CRISPR-Cas systems and other immune modules (Pinilla-Redondo et al., 2022; Grafakou et al., 2024; Samuel et al., 2024; Zheng et al., 2026). Given that MGEs are major contributors to the bacterial pan-immune system and that antimicrobial-resistance determinants frequently accumulate on plasmids, we hypothesised that plasmids may similarly serve as reservoirs for uncharacterised defence systems that shape bacterial interactions with phages. Consistent with this idea, plasmid-encoded defence systems could provide a selective advantage during phage predation while simultaneously promoting the maintenance and spread of ARG-containing plasmids (Rodríguez-Beltrán et al., 2021; Botelho et al., 2023; Beamud et al., 2024). However, many plasmid-associated defence genes remain functionally uncharacterised, and the diversity, distribution and mechanisms of these systems remain poorly understood, particularly in clinically important Gram-positive pathogens. Experimental interrogation and perturbation of MGEs therefore represent promising complementary approach for uncovering components of the bacterial pan-immune system that may escape sequence-based discovery strategies.

*Staphylococcus aureus* provides a powerful model for studying interactions between MGEs and phages. This pathogen harbours a diverse mobilome composed of plasmids, prophages, pathogenicity islands and defence-associated islands that collectively shape susceptibility to viral infection (Firth et al., 2018; Humphrey et al., 2021). Several anti-phage systems have been characterised in staphylococci, including restriction-modification systems, CRISPR-Cas systems and prophage- or satellite-associated defences (Hossain et al., 2024; Rostøl et al., 2024; Aguayo-González et al., 2026). Despite extensive characterisation of staphylococcal defence systems, comparatively little is known about the contribution of plasmids to the staphylococcal anti-phage repertoire. Given the central role of plasmids in HGT and the dissemination of adaptive traits, together with the broader contribution of MGEs to genome evolution and phage susceptibility, plasmids are likely to serve as reservoirs of previously uncharacterised defence systems that contribute to the bacterial pan-immune system and facilitate the redistribution of antiviral functions across staphylococcal populations (Bernheim and Sorek, 2020; Cui et al., 2025).

This question is particularly relevant because phage therapy has re-emerged as a strategy to treat antimicrobial-resistant infections, including infections caused by methicillin-resistant *S. aureus* (MRSA). Strictly lytic staphylococcal phages, including *Silviavirus* phages such as φMR003 and related viruses, have been proposed as antimicrobial agents against MRSA infections (Peng et al., 2019; Lerdsittikul et al., 2024). However, the success of such approaches depends on the defence landscape of the target strain. Recent work suggests that defence systems can be major determinants of host range in *S. aureus* clinical isolates and that engineered phages can be designed to evade complex MRSA defence repertoires (Voss et al., 2026). If antimicrobial-resistance plasmids encode dedicated anti-phage systems, they could couple antibiotic resistance with altered susceptibility to therapeutic phages, thereby influencing both plasmid ecology and phage-based interventions. This possibility is particularly relevant given growing interest in phage-antibiotic synergy as a strategy to combat multidrug-resistant infections. Co-selection of antimicrobial resistance and phage-defence functions on the same MGEs could complicate such approaches by simultaneously promoting antibiotic resistance and reducing phage susceptibility.

In this Article, we describe the Evangelion system, a family of plasmid-associated anti-phage systems enriched on antimicrobial-resistance plasmids and distributed across diverse bacterial hosts and MGEs. Evangelion systems are built around a conserved DUF4062-containing core linked to a variable auxiliary region and protect against phage infection in distinct host contexts. We show that Eva01, the pMw2-encoded representative, protects *S. aureus* against lytic *Silviavirus* phages, whereas related systems function in other bacterial hosts. Functional, structural and evolutionary analyses support a model in which Evangelion systems use a conserved nucleotide-metabolism-associated core that has diversified through coupling to distinct auxiliary regions. Overall, our findings identify a previously unrecognised family of mobile defence systems linking plasmid ecology, antimicrobial resistance and phage susceptibility in important pathogens.

## Results

### pMw2 encodes Eva01, a plasmid-borne defence system against *Silviavirus* phages

To investigate whether resident MGEs contribute to phage susceptibility in *S. aureus*, we systematically removed endogenous MGEs from the MRSA-Mw2 lineage and challenged the resulting strains with a panel of staphylococcal phages. Previous work established that pMw2 is stably maintained in its native host and imposes little detectable fitness cost under laboratory growth conditions (Dorado-Morales et al., 2021). Loss of the native plasmid pMw2 increased susceptibility to several *Silviavirus* phages, including φMR003, φeMR003 and φ74 (Fig. 1A). Quantification by efficiency of plating confirmed that pMw2 reduced phage propagation by several orders of magnitude, while liquid infection assays showed that pMw2-containing cultures recovered from low-MOI challenge but collapsed at higher phage burdens (Fig. 1A and Fig. S1). Thus, pMw2 confers a conditional anti-phage phenotype consistent with abortive defence.

**Figure 1.**
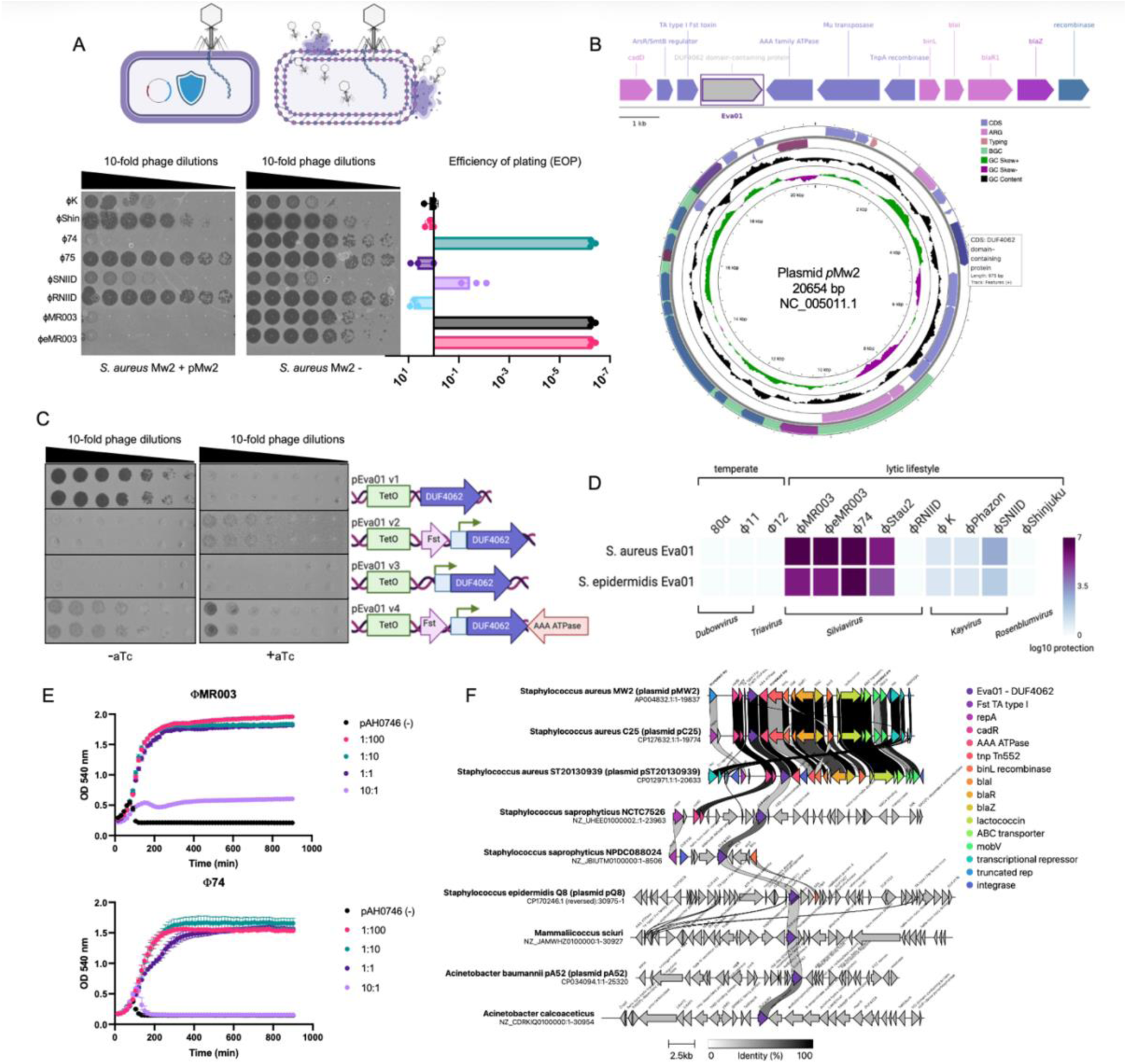
Evangelion is a plasmid-borne anti-phage system that protects Staphylococci against *Silviavirus* phages. **(A)** Loss of the native MRSA plasmid pMw2 increases susceptibility to multiple *Silviavirus* phages, as measured by spot assays, efficiency of plating (EOP) (*n*=3 ± SD)**. (B)** Genetic organisation of pMw2 showing the location of Eva01 adjacent to antimicrobial-resistance and mobility-associated genes. **(C)** Mapping of the minimal functional Eva01 locus using inducible constructs containing different DUF4062-containing regions. Representative spot assays are shown**. (D)** Defence spectrum (*n*=4) of Eva01 homologues from different bacterial hosts against a panel of staphylococcal phages. Heatmap colours indicate the magnitude of protection. **(E)** Growth curves (*n*=3) following infection with φMR003 or φ74 at the indicated phage:bacterium ratios (MOIs). **(F)** Comparative genomic organisation of representative Eva01-containing loci from diverse plasmids. Conserved Eva01 genes are highlighted and neighbouring resistance, mobility and accessory genes are shown. See also Figures S1-S4.

To identify the responsible locus, we cloned candidate plasmid regions enriched in hypothetical proteins and domains of unknown function (DUF). A single locus encoding a DUF4062-containing protein reproduced the anti-phage phenotype in both Mw2 and RN4220 (Fig. S2). Notably, this gene lies near antimicrobial resistance genes (ARGs) and mobility-associated genes (Fig. 1B), linking anti-phage defence to a clinically relevant plasmid context.

We next defined the minimal functional region, using inducible constructs carrying DUF4062 with different flanking sequences. Constructs encoding DUF4062 retained anti-phage activity following induction by anhydrotetracycline (aTc), and a minimal construct, pEva01v3, conferred robust protection even without induction, consistent with basal activity from local regulatory sequences (Fig. 1C and Fig. S3). Interestingly, the DUF4062 gene encodes a predicted two-part protein consisting of a conserved N-terminal DUF4062 region and a short C-terminal segment lacking recognised domain annotations.

We named this defence system Evangelion, inspired by the modular defensive units of Neon Genesis Evangelion. The analogy reflects both the conserved core and specialised auxiliary architecture of the system and its apparent role as an executioner-like defence that sacrifices infected cells to protect the wider bacterial population. Because this gene was sufficient to reproduce the pMw2-mediated anti-phage phenotype, we designated it Evangelion type 01 (Eva01), the founding member of a previously uncharacterised defence family.

When examining the defence spectrum, Eva01 did not confer broad phage resistance but instead displayed a selective protection profile. Strong interference was observed against *Silviavirus* phages, whereas unrelated staphylococcal phages were largely unaffected (Fig. 1D). Liquid infection experiments mirrored these results: Eva01-expressing cultures recovered following low-dose challenge with φMR003 or φ74, whereas control cultures remained cleared (Fig. 1E). These data identify Eva01 as a *bona fide* plasmid-borne anti-phage system with pronounced specificity towards lytic *Silviavirus* phages.

### Eva01 defines a widespread plasmid-associated defence lineage enriched on ARG-containing plasmids

The location of Eva01 on pMw2 raised the possibility that related defence loci are disseminated on clinically relevant plasmids. Comparative analysis revealed Eva01-like genes in diverse plasmid backbones, frequently positioned near ARGs, toxin-antitoxin modules and mobility-associated genes (Fig. 1F and Fig. S4). A closely related Eva01 homologue from *Staphylococcus epidermidis* pQ8 displayed a defence profile similar to *S. aureus* Eva01, whereas more divergent homologues from *Acinetobacter baumannii* and *Lactococcus lactis* did not protect against the staphylococcal phages tested (Fig. 1D, Fig. S5 and Table S2).

To define the broader distribution of Eva01-like systems, we screened the plasmid databases PLSDB (Molano et al., 2025) using a panel of bacterial Eva01/DUF4062 protein sequences. Non-redundant plasmids containing homologues with at least 40% query coverage per HSP were retained. This identified 447 Evangelion-positive plasmids, of which 327 carried at least one ARG (73.2%; Fig. 2A). Most were associated with *Staphylococcus* species (202 plasmids, 45.2%) or *Enterococcus* species (159 plasmids, 35.6%). Smaller numbers were identified in *Legionella* (2.9%), *Rhodococcus* (2.5%), *Acinetobacter* and *Synechococcus* (1.1%), Bacillus (0.7%) and Salmonella (0.4%) (Fig. 2B and Table S3). Notably, Eva01 homologues were detected in 202 of 3,174 Staphylococcus plasmids represented in PLSDB (6.4%), indicating that Evangelion systems occur in a substantial subset of staphylococcal plasmids.

**Figure 2.**
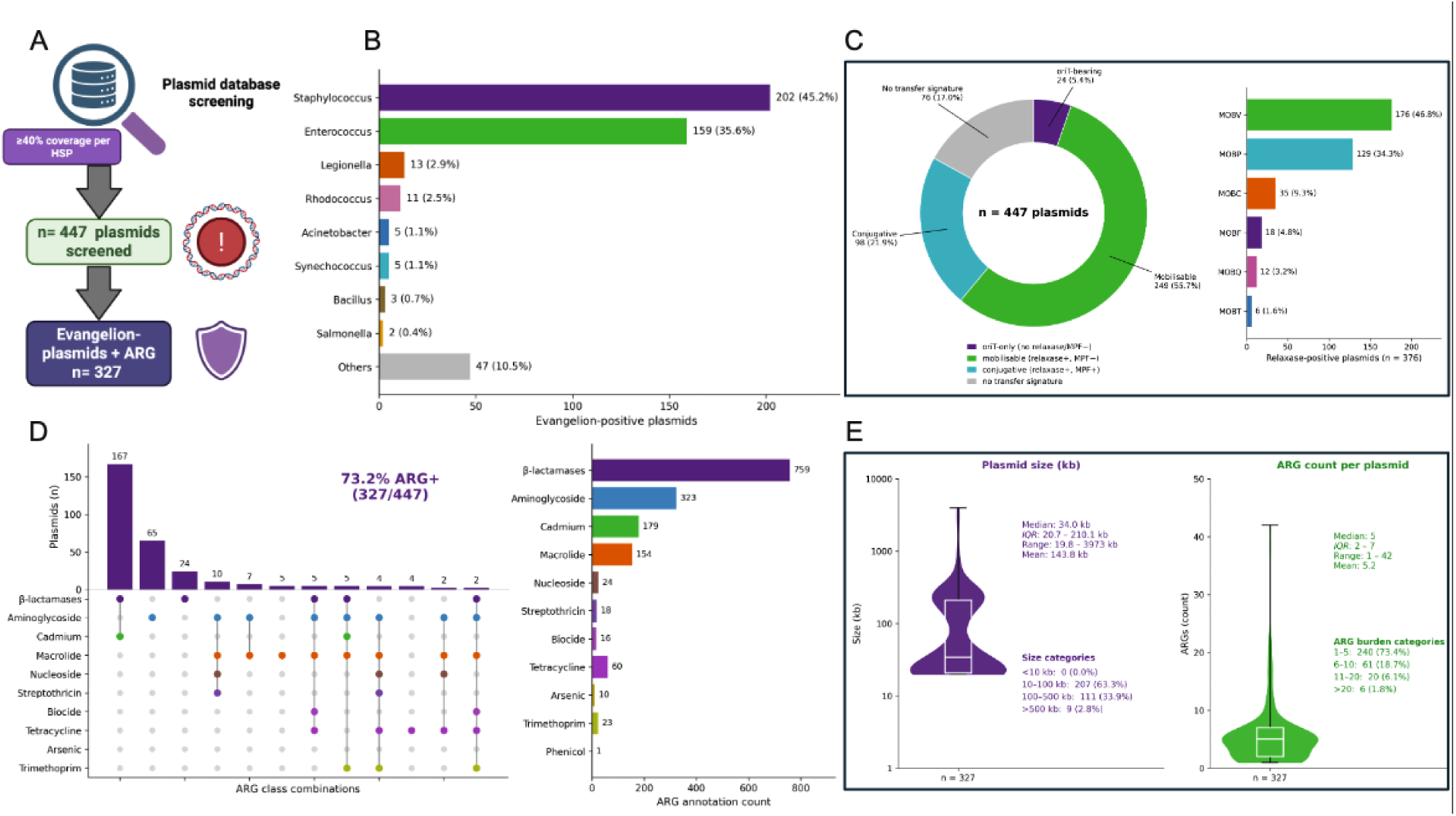
Evangelion type I (Eva01) is widespread on ARG-rich plasmids. **(A)** Workflow used to identify Eva01-positive plasmids from PLSDB and quantify associated antimicrobial-resistance genes (ARGs). **(B)** Taxonomic distribution of ARG-positive Evangelion plasmids across bacterial genera. **(C)** Distribution of transfer-associated features among ARG-positive Evangelion plasmids. Left, classification of plasmids according to predicted mobility signatures (oriT-bearing, mobilisable, conjugative, or lacking detectable transfer features). Right, distribution of relaxase families among relaxase-positive plasmids. **(D)** Most frequent ARG combinations and resistance classes identified on Evangelion-positive plasmids. **(E)** Distribution of plasmid sizes and ARG content among ARG-positive Evangelion plasmids. Violin plots indicate the distribution of plasmid sizes and ARG counts, with embedded boxplots showing medians and interquartile ranges.

We next examined transfer-associated features. Nearly half of Eva01-positive plasmids were classified as mobilisable (55.7%), 98 as conjugative (21.9%) and 24 as oriT-only but lacking an identifiable relaxase or mating-pair formation system (5.4%) (Fig. 2C). Only 76 plasmids (17%) did not have transfer signatures. Among the 376 plasmids encoding an identifiable relaxase, MOBV was the most frequent relaxase family, followed by MOBP and MOBC. These plasmids also carried diverse ARG combinations, most commonly β-lactamase, aminoglycoside, cadmium and macrolide resistance determinants (Fig. 2D). Most ranged from 10-100 kb and carried a median of five ARG annotations (Fig. 2E).

Therefore, Eva01 is not an isolated feature of staphylococcal plasmids, but a widespread plasmid-associated defence lineage enriched on ARG-rich and frequently transferable plasmids.

### Evangelion systems encode a conserved DUF4062/TIR-like effector core

Having established Eva01 as the founding member of a widespread plasmid-associated defence lineage, we next defined the molecular architecture of Evangelion proteins. Conserved domain analysis (Wang J et al, 2023) and AlphaFold3 (Abramson et. al, 2024) predicted that Eva01 contains a compact N-terminal DUF4062 region with an α/β fold resembling TIR-like proteins, followed by a short C-terminal auxiliary region (AUX) (Fig. 3A). A putative ADP/NAD⁺-binding pocket was identified within the DUF4062/predicted TIR-like core, suggesting that this region may contribute to nucleotide processing.

**Figure 3.**
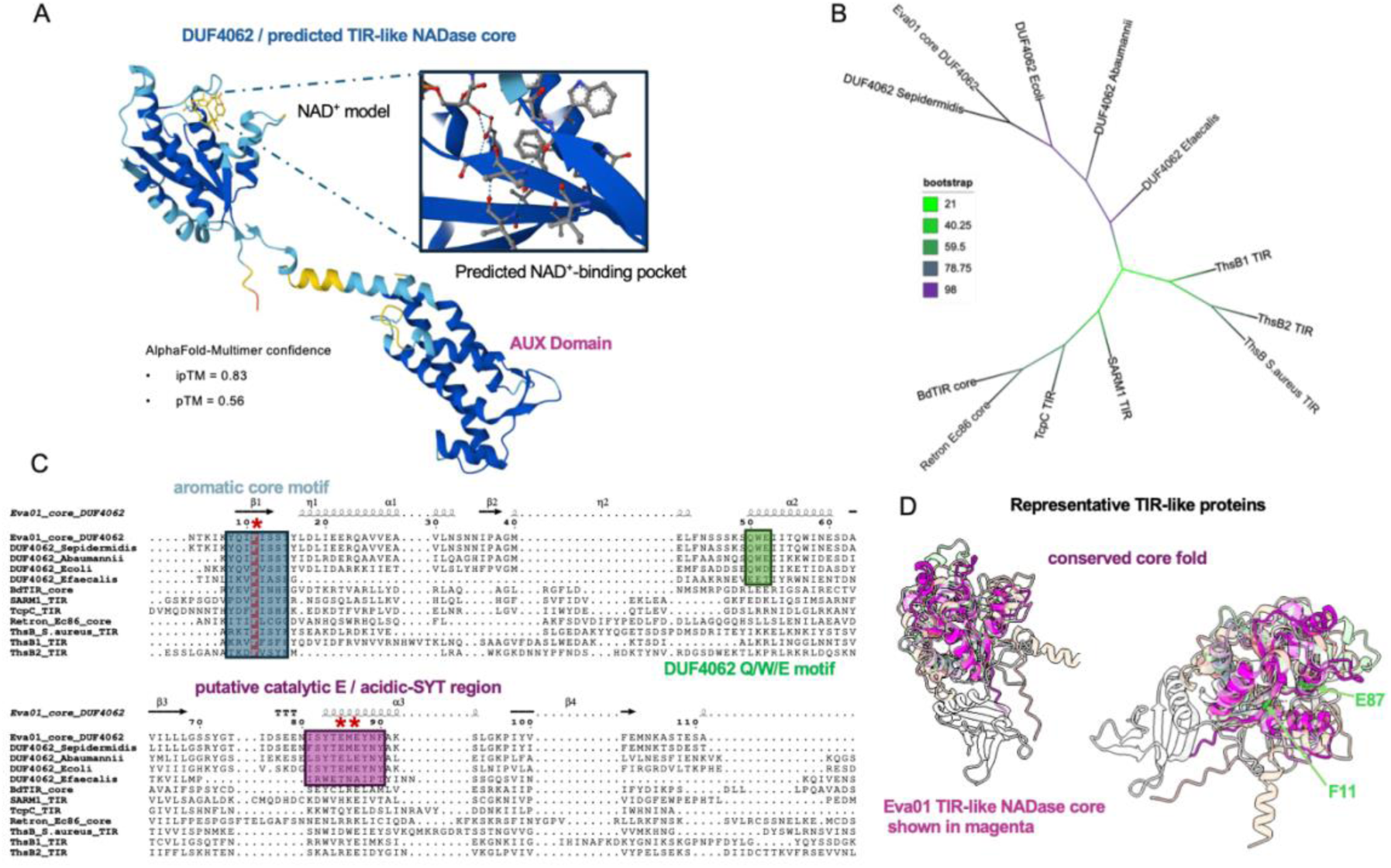
Evangelions contain a divergent DUF4062/TIR-like NADase core with conserved catalytic motifs. **(A)** AlphaFold-Multimer model and domain organization of representative Eva01 proteins. The N-terminal DUF4062/predicted TIR-like NADase core is shown with a zoomed view of a predicted NAD⁺-binding pocket**. (B)** Maximum-likelihood tree of selected Eva/DUF4062 cores and representative TIR-like proteins. Branch colours indicate bootstrap support. **(C)** Multiple sequence alignment of selected Eva/DUF4062 cores and representative TIR-like proteins. Highlighted regions indicate a conserved aromatic core motif, a DUF4062 Q/W/E motif, and an acidic-SYT region containing the putative catalytic Glu. Red asterisks mark residues selected for mutagenesis. **(D)** Structural superposition of Eva01 with representative TIR-like proteins, showing conservation of the core fold. Eva01 is shown in magenta, with F11 and E87 indicated. See also Figures S5-6.

Although DUF4062-containing proteins have been identified in diverse bacterial genomes, their biological function remains poorly understood (Wang et al., 2023). To place Evangelion proteins in an evolutionary context, we compared representative Eva/DUF4062 cores with bacterial and eukaryotic TIR domains and TIR-like proteins, including ThsB-family sensors and nucleotide-associated defence proteins (Essuman et al., 2017; Horsefield et al., 2019; Wan et al., 2019; Ofir et al., 2021; Leavitt et al., 2022; Ka et al., 2020; Roberts et al., 2025). Maximum-likelihood phylogenetic analysis resolved Evangelion proteins as a distinct DUF4062-associated lineage that was clearly separated from canonical bacterial TIR proteins and ThsB-family sensors (Fig. 3B). These results suggest that Evangelion proteins belong to a broader group of TIR-like nucleotide-associated proteins (Wang et al., 2023; Osinski et al., 2026).

To examine conservation within the family, we aligned representative Evangelion proteins spanning diverse bacterial hosts and MGE contexts. Despite extensive divergence within the AUX region, the DUF4062 core contained several highly conserved motifs (Fig. 3C and Fig. S6). These included an N-terminal aromatic motif centred on phenylalanine F11, a conserved acidic-SYT region containing a putative catalytic glutamate E87, and a DUF4062-specific region encompassing glutamine-tryptophan-glutamic acid Q/W/E. Conservation of these residues across phylogenetically distant Evangelion proteins suggested that they contribute to a shared biochemical function.

Using AlphaFold3 modelling and structural superposition with representative TIR and TIR-like proteins, we found a conserved fold of the DUF4062 core despite limited primary-sequence similarity (Fig. 3D). Mapping conserved residues onto the predicted model positioned F11 and E87-Y88 within a common surface pocket of the DUF4062 core, consistent with a potential catalytic or ligand-binding role.

### The DUF4062 core and AUX region function cooperatively during Evangelion defence

The predicted DUF4062 pocket and the conservation of F11, E87 and W162 suggested that the core region of Eva01 may provide the biochemical output of the system. To test this, we generated an evolution-guided panel of Eva01 variants targeting the DUF4062 core, the DUF4062-AUX junction and the AUX region (Fig. 4A-B and Fig. S5). Because lysine/arginine-density analysis revealed positively charged patches near the DUF4062-AUX interface and within AUX, we also included mutations designed to probe regions with potential nucleic-acid-interaction properties (Fig. 4B).

**Figure 4.**
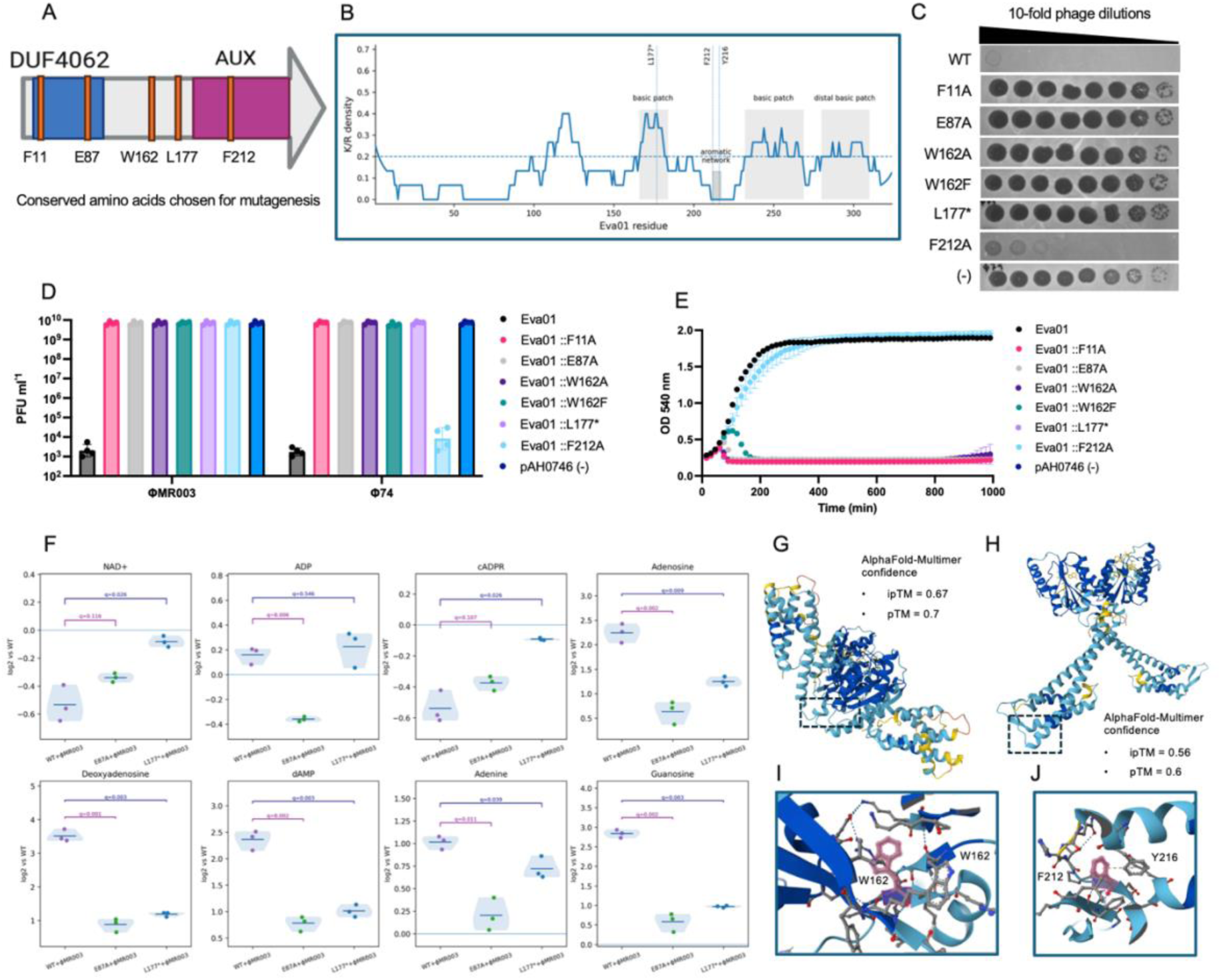
Anti-phage defence requires cooperation between the DUF4062 core and AUX region. **(A)** Schematic of Eva01 variants targeting conserved DUF4062 residues, the DUF4062-AUX interface and the AUX region. **(B)** Evolutionary conservation and residue-selection strategy used for mutagenesis. Conserved residues selected for functional analysis are indicated. **(C)** Representative spot assays showing the effect of Eva01 mutations on defence against *Silviavirus* phages. **(D)** Quantification of phage propagation (*n*=4 ± SD) following infection with φMR003 and φ74. **(E)** Liquid infection assays (*n*=3) demonstrating the effects of DUF4062 and AUX mutations on protection against φMR003. **(F)** Untargeted LC-MS analysis (*n*=3 ± SD) of selected adenine-associated metabolites following φMR003 infection in strains expressing wild-type or mutant Eva01 variants. **(G-H)** AlphaFold-Multimer models of Eva01 oligomeric assemblies. **(I-J)** Structural views highlighting conserved residues W162 and F212 within the predicted DUF4062-AUX interface and associated pockets. See also Figure S7.

Spot assays and phage-yield measurements showed that F11A, E87A, W162A, W162F and the AUX truncation Eva01::L177* disrupted defence against *Silviavirus* phages, producing susceptibility and phage titres comparable to inactive controls (Fig. 4C,D). W162F retained only weak residual activity in liquid infection assays, indicating that aromatic substitution is insufficient to replace the native tryptophan at this position (Fig 4E). By contrast, Eva01::F212A retained protection against φ74 but showed impaired defence against φMR003, suggesting that individual AUX residues can differentially affect responses to distinct *Silviavirus* phages.

Because structural analyses identified a putative nucleotide-associated pocket within DUF4062, we next examined the metabolic consequences of Evangelion activation during phage infection. Untargeted polar LC-MS analysis performed 15 minutes after φMR003 challenge revealed pronounced alterations in adenine-associated metabolites in cells expressing wild-type Eva01 relative to inactive mutants (Fig. 4F). Compared with E87A and L177*, wild-type Eva01 displayed reduced NAD⁺ abundance together with broader perturbations in adenine-containing metabolites, including, guanosine adenosine, adenine and related nucleotide derivatives. Although these measurements do not directly establish enzymatic activity, they indicate that Evangelion activation is associated with nucleotide-metabolic remodelling during infection (Fig. 4F and Fig. S7).

To investigate how the DUF4062 core and AUX region might communicate, we modelled Eva01 using AlphaFold-Multimer. The resulting structure positioned the DUF4062 domain and AUX region in close physical proximity despite their distinct folds, with W162 near the predicted dimer interface and F212 within a conserved AUX pocket (Fig. 4G-J). Modelling of the N-terminal DUF4062-containing region further predicted dimeric assemblies with stacked interfaces in both NAD⁺- and ADP-bound models (Fig. S8A-B). Consistent with this, recombinant Eva01 purified from *E. coli* resolved by native mass spectrometry as multiple species, including masses compatible with dimeric and higher-order assemblies (Fig. S8C). These observations suggest that Eva01 can self-associate, although the role of oligomerisation in defence activation remains to be determined.

### Phage escapers implicate replication proteins in Evangelion activation

To identify candidate determinants of Evangelion susceptibility, we isolated spontaneous escaper mutants of φMR003 and φ74 capable of infecting Eva01-containing cells (Fig. 5A and S9A-B). Whole-genome sequencing revealed that independent φMR003 escapers repeatedly acquired mutations in a region encoding a RecA/Sak-like recombination protein and a neighbouring homing-endonuclease-remnant gene (HEG) (Fig. 5B). Similarly, φ74 escapers accumulated mutations within a UvsX-like recombination protein and an adjacent hypothetical gene (Fig. 5C).

**Figure 5.**
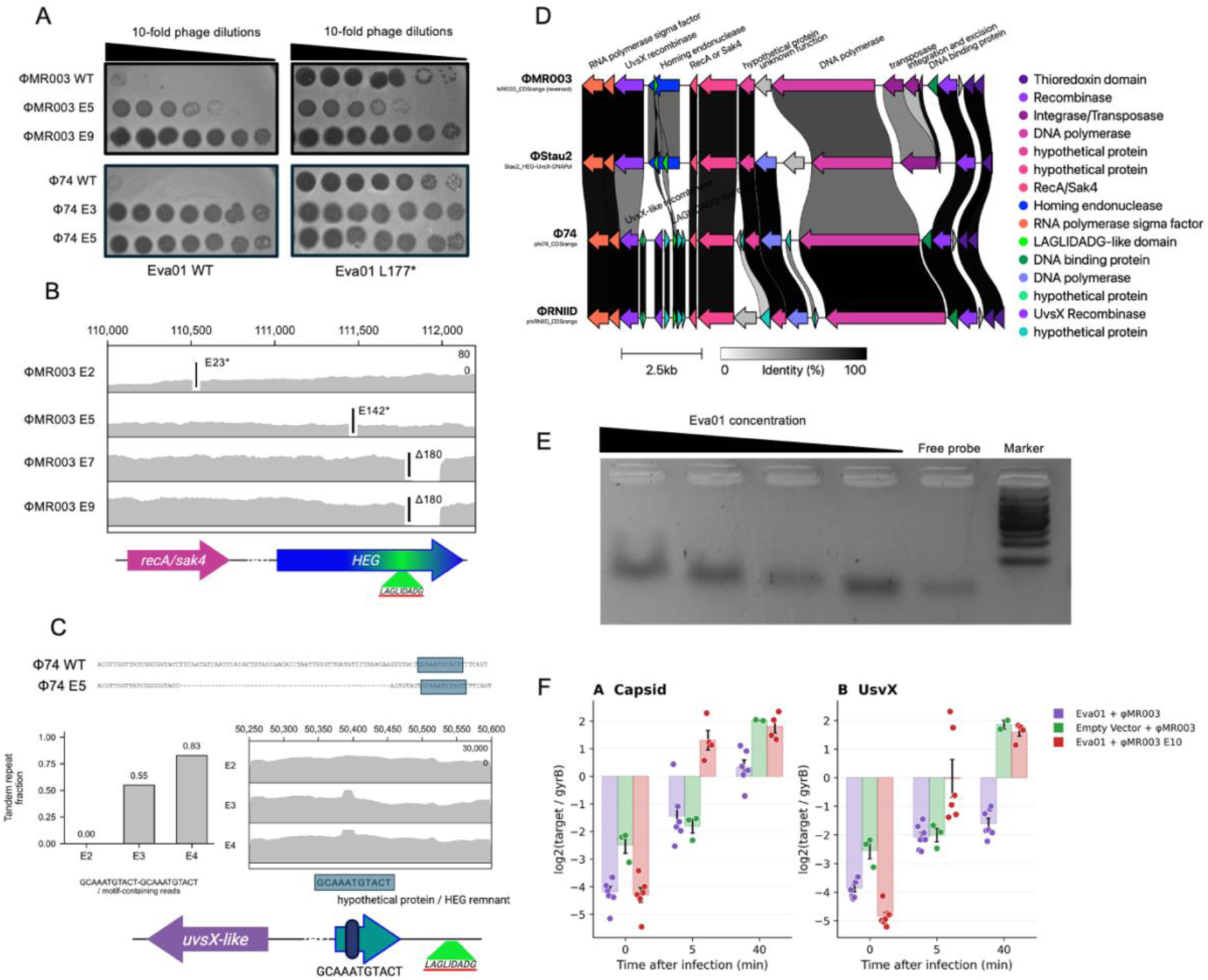
Phage escapers reveal replication-associated triggers of Evangelion defence. **(A)** Isolation of spontaneous φMR003 and φ74 escaper mutants capable of infecting Eva01-expressing cells. **(B)** Genomic locations of mutations identified in independent φMR003 escapers. **(C)** Genomic locations of mutations identified in independent φ74 escapers. **(D)** Comparative analysis of *Silviavirus* genomes showing that escaper mutations converge on a conserved module enriched in recombination- and replication-associated genes, including RecA/Sak-like and UvsX-like proteins. **(E)** Eva01 preferentially associates with recombination-like DNA substrates relative to fully duplex DNA. Representative EMSA assays are shown. **(F)** Digital PCR quantification of intracellular phage DNA following infection with wild-type φMR003 or the escaper mutant φMR003 E10. Phage capsid and *usv*X copy numbers were normalized to the chromosomal reference gene *gyr*B and plotted as log2(target/*gyr*B). Bars indicate the mean ± SD, and points *n ≥* 3 independent biological replicates per condition. See also Figures S8-9.

To determine whether these loci were shared among Eva01-sensitive phages, we compared the corresponding genomic regions across multiple *Silviavirus* genomes. Comparative analysis showed that these loci reside within a conserved module enriched in recombination- and replication-associated genes, including RecA/Sak-like and UvsX-like homologues (Fig. 5D and S9C). The convergence of escaper mutations on this shared module suggested that Eva01 activation is associated with conserved replication functions rather than highly variable structural components of the virion.

### Eva01 restricts intracellular phage genome accumulation after DNA entry

Because escapers repeatedly targeted phage recombination-associated proteins, we asked whether Eva01 could interact with DNA substrates that mimic replication or recombination intermediates. Purified Eva01 was incubated with double-stranded DNA substrates containing single-stranded overhangs designed to resemble structures generated during homologous recombination. Under these conditions, Eva01 associated preferentially with recombination-like substrates relative to fully duplex DNA (Fig. 5E). This substrate preference is consistent with the genetic evidence implicating replication-associated phage functions in Evangelion susceptibility.

We next tested whether Eva01 affects phage genome accumulation during infection. Cultures carrying Eva01 or an empty-vector control were challenged with wild-type φMR003, whereas Eva01-containing cells were additionally infected with the escaper mutant φMR003 E10. Intracellular phage DNA abundance was quantified by digital PCR using primers targeting two independent genomic loci and normalised to the *S. aureus* reference gene *gyr*B. Digital PCR targeting the capsid and *uvs*X genes showed that both phage markers remained detectable following infection, demonstrating that phage DNA enters Eva01-containing cells (Fig. 5F). However, accumulation of both loci was markedly reduced relative to the empty-vector control, indicating that Eva01 acts after DNA injection and restricts productive phage genome amplification. Early increases in phage DNA abundance further suggest that replication is initiated but does not proceed efficiently in the presence of Eva01.

In contrast, φMR003 E10 restored intracellular phage DNA accumulation despite the presence of Eva01 (Fig. 5F). Similar trends were observed for both capsid and *uvs*X, indicating that the effect was not restricted to a specific region of the phage genome. Thus, escape mutations that enable plaque formation in Eva01-containing cells also restore phage genome accumulation during infection.

Although the precise activating signal remains unknown, the repeated selection of mutations within homologous recombination-associated loci across independent escapers and distinct *Silviavirus* phages points to a common underlying mechanism. The convergence of mutations in UvsX-like and RecA/Sak-like functions, together with the observation that phage genomes are detectable in Eva01-containing cells but fail to accumulate efficiently thereafter, suggests that Evangelion defence is linked to conserved replication-associated processes. Whether Eva01 directly senses phage recombination proteins or instead responds to replication intermediates generated during infection remains an important question for future investigation.

### DUF4062-containing proteins form evolutionarily distinct defence-associated lineages

Having established Eva01 as the founding member of the Evangelion defence family, we next investigated the evolutionary diversity of DUF4062-containing proteins in bacteria. We analysed approximately 6,000 bacterial proteins containing the DUF4062 domain (PF13271) and used this dataset to examine the distribution of DUF4062 containing proteins across bacterial genomes and MGEs. Phylogenetic analysis resolved several distinct DUF4062-associated lineages recovered from plasmids, prophages, phage satellites, integrative elements and chromosomal loci (Fig. 6A-B and S10A). Among these, Eva01 formed a discrete clade together with several related DUF4062 proteins that we collectively refer to as Evangelion lineages, separated from additional groups represented by Eva02, Eva03, Eva04, the Swarożyc system and bacterial TEP1-like proteins.

**Figure 6.**
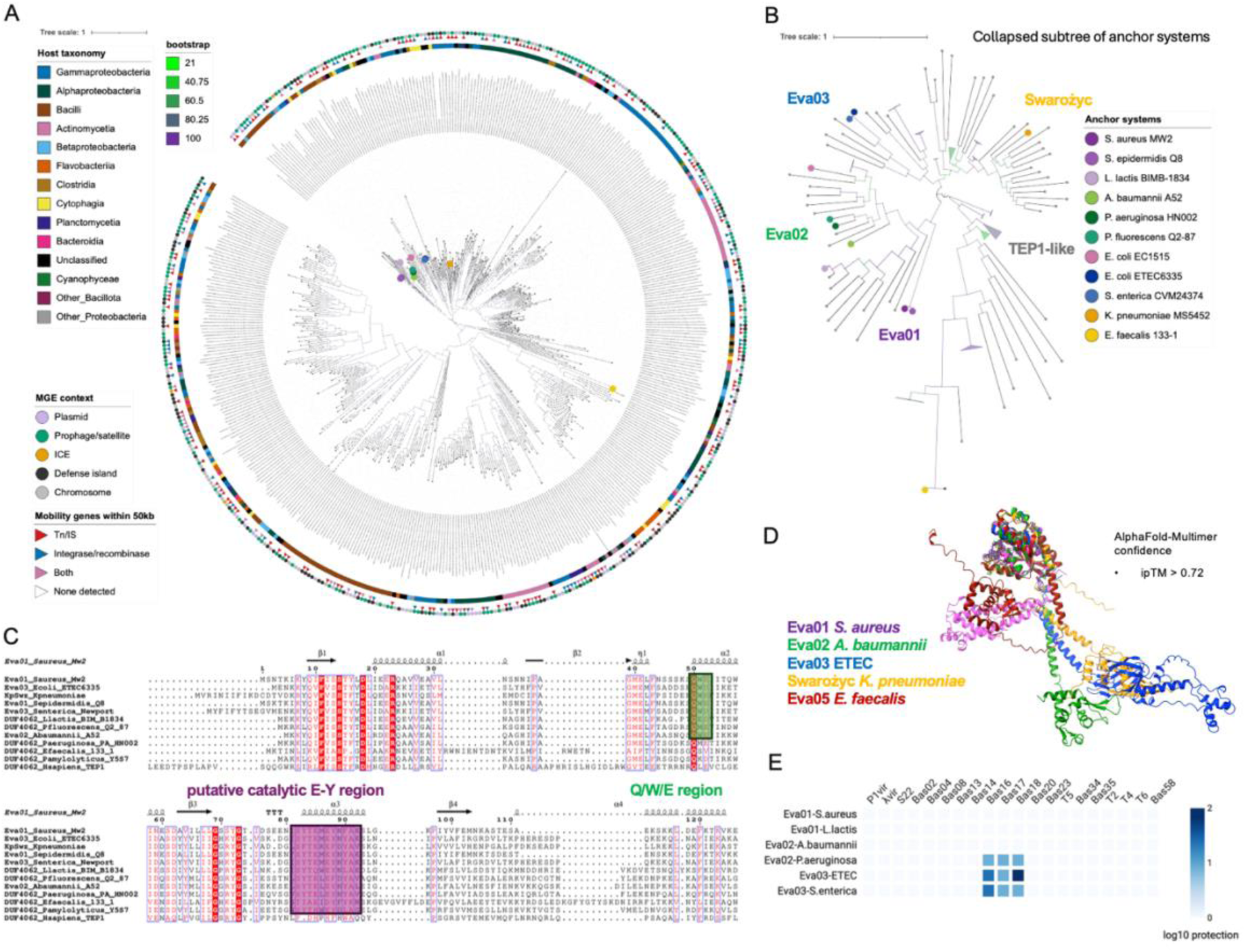
Evangelion proteins form diverse DUF4062 defence lineages with distinct antiviral activities. **(A)** Phylogenetic distribution of approximately 6,000 DUF4062-containing proteins across bacterial genomes and MGEs. Metadata tracks indicate taxonomic origin and genomic context. **(B)** Expanded view of major Evangelion-associated lineages, including Eva01, Eva02, Eva03, Swarożyc and TEP1-like proteins. **(C)** Alignment of representative Evangelion proteins highlighting conservation of the putative catalytic E-Y region and DUF4062 Q/W/E motif. **(D)** Structural comparison of representative Evangelion proteins from diverse bacterial hosts showing conservation of the DUF4062 core despite divergence of auxiliary regions. **(E)** Defence profiles of representative Evangelion systems against host-specific phage panels. Heatmap colours indicate the magnitude of protection. See also Figure S10.

Despite substantial sequence divergence, representative proteins retained several conserved DUF4062 motifs, including the putative catalytic E-Y region and the characteristic Q/W/E motif (Fig. 6C). Structural modelling further supported a common evolutionary origin. AlphaFold predictions of representative Evangelion proteins from *S. aureus*, *A. baumannii*, *Escherichia coli*, *Enterococcus faecalis* and the Swarożyc system of *Klebsiella pneumoniae* revealed a conserved DUF4062 core architecture despite extensive divergence elsewhere in the protein (Fig. 6D and S10B). In contrast, the auxiliary regions varied markedly in length, topology and predicted surface properties. Consistent with this broader relationship, CD searches identified DUF4062 as the shared feature between Eva01 and bacterial TEP1-like proteins, whereas TEP1-like homologues contained additional TROVE, NACHT and WD40-repeat domains, consistent with distinct biological contexts (Fig. S11). The presence of these domains strongly supports an evolutionary link to eukaryotic TEP-1 proteins of the RNA vault.

These observations indicate that Evangelion proteins belong to a broader and evolutionarily diverse superfamily of DUF4062-containing proteins that share a conserved structural core.

### Distinct Evangelion lineages exhibit specialised defence profiles

The conservation of the DUF4062 core together with extensive diversification of auxiliary regions suggested that distinct Evangelion lineages may have evolved specialised defence activities. To test this possibility, representative systems from different lineages were challenged with phage panels in their respective experimental hosts.

Interestingly, Eva01 preferentially restricted staphylococcal *Silviavirus* phages, including φMR003 and φ74 (Fig. 6E and Table S2). By contrast, Eva03 displayed a distinct defence profile and interfered with members of the BASEL phage collection belonging to the *Dhillonvirus* genus. Although the magnitude of protection conferred by Eva03 was lower than that observed for Eva01, these experiments demonstrate anti-phage activity in a phylogenetically distant bacterial host. In contrast, Eva02 did not confer detectable protection against the phages tested in either *S. aureus* or *E. coli* under the conditions examined.

Collectively, these findings define Evangelion as a broader family of DUF4062-based defence systems that has diversified across bacterial hosts and MGEs while retaining a conserved structural core. The distinct defence spectra of Eva01 and Eva03 further indicate that diversification of Evangelion proteins is accompanied by diversification of anti-phage activity. Thus, conservation of the DUF4062 core is compatible with extensive functional diversification, allowing distinct Evangelion lineages to adapt to different host-phage environments.

### DUF4062 and AUX regions function as co-evolved defence modules

The conservation of DUF4062 across Evangelion lineages and bacterial TEP1-like proteins raised the question of whether this domain retains functional features across distant biological contexts. Structural comparison of Eva01 and human TEP1 revealed similarity between their DUF4062 regions, including a predicted nucleotide-binding pocket within the shared core fold (Fig. 7A and S12). In TEP1, DUF4062 is embedded within a large multidomain protein, whereas in Eva01 it is coupled to a compact AUX region that is required for anti-phage defence (Fig. 7B). To test whether the eukaryotic DUF4062 domain could function within an Evangelion protein, we generated chimeras in which segments of the Eva01 DUF4062 region were progressively replaced with the corresponding region from human TEP1 while retaining the Eva01 AUX domain (Fig. 7C). Complete replacement of the Eva01 DUF4062 region with the corresponding TEP1 segment failed to restore defence (Fig. 7D-E). However, several intermediate chimeras retained weak but reproducible activity against φMR003 and φ74, most notably Eva01::HsTEP1-E. Although substantially weaker than wild-type Eva01, these constructs consistently reduced phage susceptibility relative to inactive controls, indicating partial functional compatibility between bacterial and eukaryotic DUF4062 domains.

**Figure 7.**
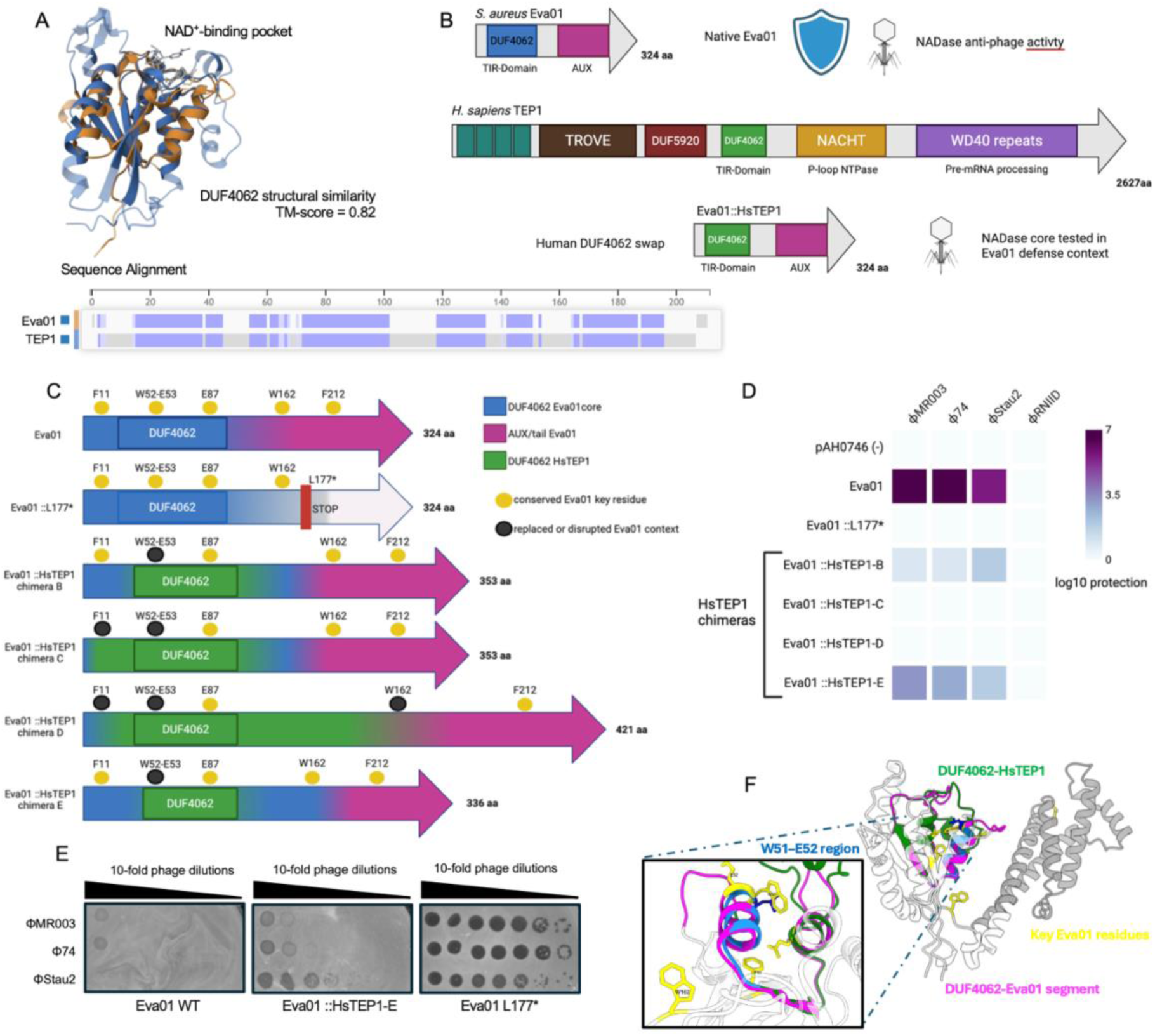
Human TEP1 DUF4062 fragments partially substitute for the Evangelion core. **(A)** Structural comparison and sequence alignment of Eva01 and the DUF4062/TIR-like region of human TEP1. **(B)** Design rationale for replacing Eva01 DUF4062 segments with the corresponding HsTEP1 region while retaining the Eva01 AUX/tail. **(C)** Schematic of Eva01::HsTEP1 chimeras. Yellow circles indicate conserved/key Eva01 residues or motifs, whereas black circles indicate regions in which the native Eva01 context is replaced or altered. **(D)** Heatmap of phage protection by Eva01, Eva01 L177*, and HsTEP1 chimeras. **(E)** Representative spot assays showing strong Eva01 defence, partial activity of Eva01::HsTEP1-E, and loss of defence in Eva01 L177*. **(F)** AlphaFold model overlay of Eva01 WT and Eva01::HsTEP1-E. The native Eva01 DUF4062 segment is shown in magenta, the HsTEP1-derived segment in green, and key Eva01 residues in yellow. The E chimera preserves the overall core architecture but alters the local W51-E52/Q-W-E region, consistent with partial rather than complete functional rescue. See also Figures S11-12.

**Figure 8.**
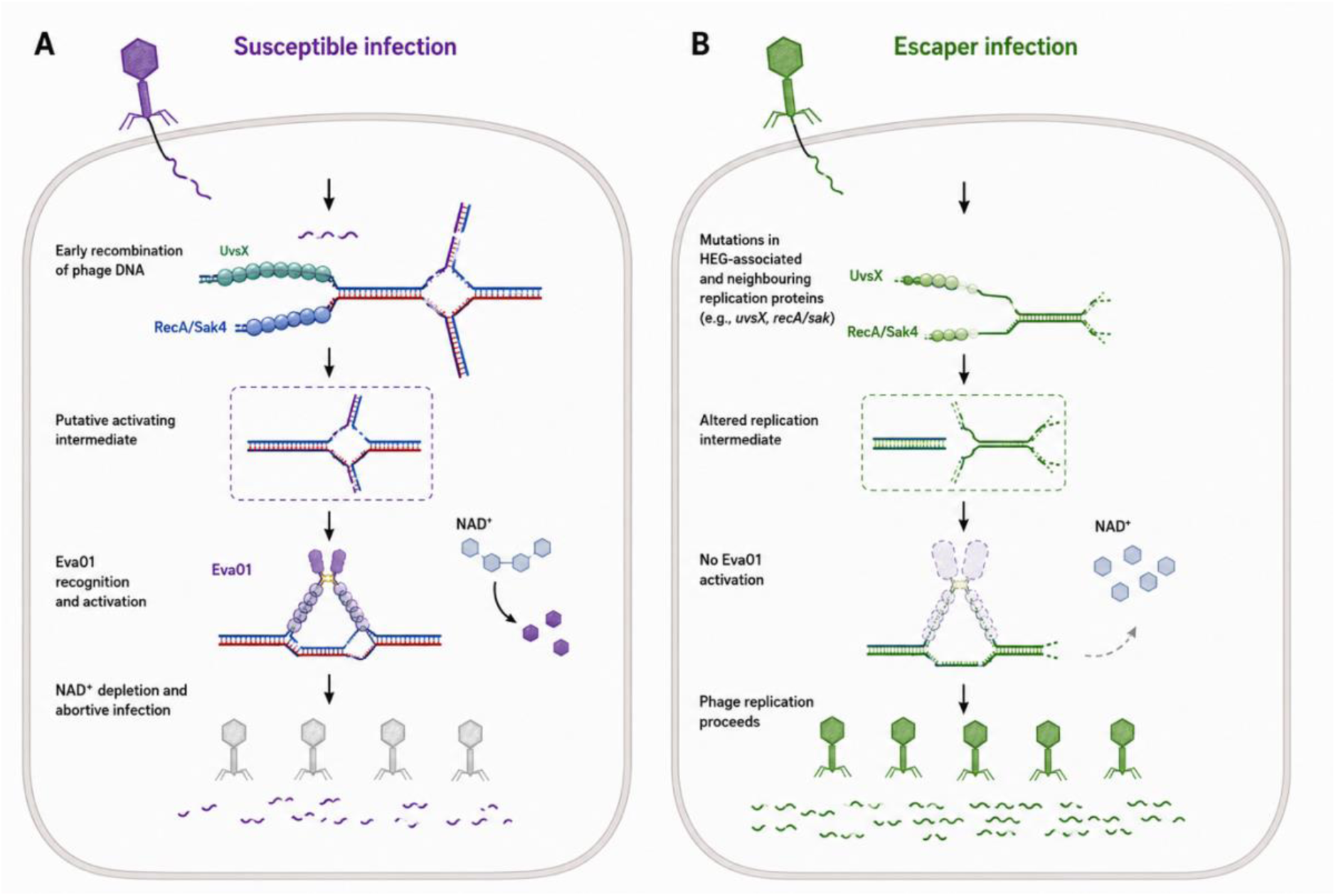
A model for Evangelion type 1 sensing of replication-associated phage intermediates. **(A)** During susceptible infection, phage recombination and replication generate UvsX- and RecA/Sak4-associated DNA structures that are recognised by Eva01. Activation of the DUF4062 defence protein triggers NAD⁺ depletion and abortive infection, restricting productive phage replication. **(B)** Escaper mutations in HEG-associated functions and neighbouring replication genes alter the activating intermediate, preventing Eva01 activation. As a consequence, NAD⁺ levels are maintained and productive phage replication proceeds.

Structural modelling of the chimeric proteins showed that active and inactive chimeras retained similar overall architectures, arguing against gross misfolding as the sole explanation for loss of defence (Fig. 7F). Instead, the strongest residual activity was observed in constructs preserving regions surrounding conserved Eva01 residues and the predicted DUF4062-AUX interface. This pattern suggests that defence requires local compatibility between the DUF4062 core and the Eva01 AUX region.

The partial activity of TEP1-derived chimeras is notable given recent work showing that human TEP1 functions as a DUF4062-dependent NADase within vault ribonucleoprotein particles (Osinski et al., 2026). These results suggest that DUF4062 domains retain conserved biochemical features across distant biological contexts, but that robust anti-phage defence requires a co-adapted DUF4062-AUX unit.

## Discussion

An intriguing finding of this study is that antimicrobial-resistance plasmids can encode previously unrecognised anti-phage systems rooted in the DUF4062 protein family. Eva01 was discovered on the MRSA plasmid pMw2, where it protects against lytic *Silviavirus* phages and forms part of a broader family of mobile defence systems distributed across diverse bacterial hosts and MGEs. Thus, pMw2 illustrates how clinically relevant plasmids are not merely vehicles for antimicrobial resistance, but also important reservoirs of antiviral immunity. This observation is consistent with recent studies identifying diverse defence systems on plasmids (Grafakou et al., 2024; Zheng et al., 2026) and supports the emerging view that MGEs contribute substantially to the bacterial pan-immune system (Bernheim and Sorek, 2020)

The strong association of Eva01-like systems with ARG-rich plasmids suggests that phage defence may be one of the accessory traits that shapes the success of resistance plasmids. Plasmid persistence is usually considered through the balance between mobility, fitness cost, antibiotic selection and host range (Frost et al., 2005; Smillie et al., 2010; San Millan, 2018; Partridge et al., 2018). Our findings add phage predation to this framework. By protecting host populations during viral attack, plasmid-encoded defence systems could provide episodic but strong selective benefits that favour maintenance of ARG-containing plasmids even outside antibiotic exposure. More broadly, recent work has shown that plasmid evolution is shaped by population-level interactions among MGEs, antibiotics and phages, with ecological selection determining which plasmid architectures are maintained over time (Rodríguez-Beltrán et al., 2021; Sastre-Dominguez et al., 2026; Ipoutcha et al., 2026). In addition, conjugative plasmids encode anti-defence systems that promote establishment during HGT, further illustrating how plasmid cargo can overcome immunity and shape dissemination (Samuel et al., 2024). In this context, phage defence systems such as Evangelion may provide an additional selective advantage that promotes the maintenance and dissemination of resistance plasmids even in the absence of direct antibiotic selection. Conversely, the same defence cargo could influence plasmid ecology by altering phage-mediated transfer, host survival and competition among co-resident MGEs.

In *S. aureus*, plasmids, prophages, pathogenicity islands and defence-related islands collectively shape genome evolution and phage susceptibility (Humphrey et al., 2021; Hossain et al., 2024; Kuang et al., 2026; Voss et al., 2026). Our study suggests that resistance plasmids can also encode potent anti-phage systems. Consequently, the MGE content of a strain may represent an underappreciated determinant of therapeutic phage efficacy, particularly for MRSA and other ESKAPE pathogens.

Our data identify DUF4062 as the conserved core of Evangelion systems. DUF4062-containing proteins are widespread but remain poorly characterised, and our structural analyses place Evangelion proteins within a divergent TIR-like family. This finding is consistent with the growing appreciation that TIR-domain and TIR-like proteins function as nucleotide-processing effectors in bacterial immunity (Ofir et al., 2021; Wang et al., 2024). In Eva01, conserved DUF4062 residues were required for defence, and phage infection was associated with pronounced alterations in NAD⁺-linked and adenine-associated metabolites. Although these observations do not establish the biochemical activity of Eva01 directly, they support a connection between Evangelion activation and nucleotide metabolism.

Notably, TIR-like domains are not universally associated with antiviral defence. The *S. aureus* TIR-domain protein TirS acts as a virulence factor that suppresses host Toll-like receptor signalling and promotes bacterial persistence during infection rather than mediating bacterial immunity (Askarian et al., 2014). More broadly, TIR and TIR-like proteins have diversified extensively across bacteria, plants and animals, where they participate in functions ranging from immune signalling to host-pathogen interactions and stress responses (Lapin et al., 2022; Yirmiya et al., 2025). The evolutionary relationship between Evangelion proteins and TEP1-like DUF4062 proteins therefore raises the possibility that DUF4062 proteins have undergone similar functional diversification while the domain itself has retained conserved biochemical properties.

The recent demonstration that a DUF4062-containing protein can function as TIR-like NADases provides a useful framework for interpreting these results. In particular, Osinski and colleagues reported that human TEP1 functions as an NADase within vault ribonucleoprotein particles (Osinski et al., 2026). Although the biological contexts differ substantially, the partial functionality of TEP1-derived chimeras in the Eva01 background suggests that bacterial Evangelion systems and eukaryotic TEP1-like proteins retain features of a common biochemical scaffold. Whether DUF4062 proteins share a conserved catalytic mechanism remains unresolved.

Our mutational and chimera analyses further demonstrate that Evangelion activity depends on cooperation between the conserved DUF4062 core and the variable AUX region. AUX truncation abolished defence, mutations at the DUF4062-AUX interface impaired activity, and replacement of Eva01 DUF4062 segments with human TEP1 sequences restored only partial protection despite preserving overall protein architecture. These observations support a model in which DUF4062 and AUX regions function as co-evolved modules rather than independently interchangeable parts. Similar architectures are common among bacterial immune systems, where conserved effector domains are coupled to variable sensing or regulatory modules that determine activation specificity (Doron et al., 2018; Payne et al., 2021; Rousset et al., 2022).

The escaper and dPCR experiments place Evangelion activity after phage DNA entry. Independent escaper mutants repeatedly acquired mutations within a conserved genomic region associated with HEG-related functions and neighbouring RecA/Sak-like and UvsX-like proteins involved in phage recombination and replication. These mutations arose independently in two distinct *Silviavirus* phages and converged on a shared replication-associated module, suggesting that Eva01 activation is connected to conserved intracellular processes rather than phage-specific structural components. Consistent with this model, Eva01 preferentially associated with recombination-like DNA substrates and reduced intracellular phage genome accumulation without preventing DNA entry. Together, these observations place Evangelion among defence systems that act after genome injection and suggest that activation occurs during early stages of phage genome processing. Whether Eva01 responds directly to phage recombination proteins, recognises replication-associated DNA structures, senses HEG-associated activities or detects downstream metabolic perturbations remains unresolved. The repeated recovery of escaper mutations within a conserved HEG-associated module, together with recent evidence that homing endonucleases can promote anti-defence functions during infection (Chihara et al., 2026), identifies this region as a promising target for future mechanistic studies.

The relatively narrow defence spectrum of Eva01 contrasts with the broader activity reported for the DUF4062 system Swarożyc (KpSwz) against the BASEL phage collection (Osinski et al., 2026). This difference may reflect distinct trigger-recognition strategies within the broader DUF4062 family. In KpSwz, phage escape mutations map to portal proteins and the AUX region engages phage structural components to activate DUF4062-dependent NADase activity. By contrast, Eva01 escapers repeatedly map to RecA/Sak-like and UvsX-like loci associated with phage DNA metabolism, and Eva01 restricts phage genome accumulation after DNA entry. Thus, although KpSwz and Evangelion proteins may share a conserved DUF4062 effector logic, their activation pathways appear fundamentally different. Notably, both systems require auxiliary (AUX) regions that are essential for defence yet lack recognizable conserved domains. In KpSwz, the AUX region functions as a phage sensor that detects portal-associated structural proteins, whereas our genetic and functional analyses implicate replication-associated processes in Eva01 activation. Together with the extensive sequence diversification observed across Evangelion lineages, these findings suggest that AUX regions may commonly function as sensory modules that determine trigger recognition, while the DUF4062 core provides a conserved defence effector. Portal-associated sensing could permit broad recognition of phages carrying compatible structural determinants, whereas replication-linked sensing may impose stricter requirements on the timing, structure or protein composition of the infection intermediate. This distinction may explain why Eva01 displays pronounced host- and phage-specific activity despite retaining a conserved DUF4062 core.

The broader evolutionary distribution of Evangelion systems supports the idea that bacterial immunity evolves within a highly mobile genetic landscape (Ibarra-Chávez et al., 2022). Recent computational and experimental surveys indicate that currently known defence systems likely represent only a fraction of the bacterial pan-immune repertoire and that many defence-associated protein families remain functionally uncharacterised (Bernheim and Sorek, 2020; Mordret et al., 2026). Defence systems are pervasive across plasmids, prophages, phage satellites and other MGEs, reflecting ongoing conflicts among phages, bacterial hosts and competing genetic elements (Botelho et al., 2023; Beamud et al., 2024; Pfeifer et al., 2022). In this context, Evangelion appears to represent a mobile defence module that has diversified across distinct evolutionary and ecological settings. Notably, unlike many bacterial immune systems that require multiple proteins for sensing, signalling and interference, Evangelion is encoded by a single gene that combines a conserved DUF4062 effector core with a highly variable auxiliary region. This compact architecture may facilitate HGT and stable maintenance on plasmids and other MGEs, while allowing diversification of trigger-recognition functions through AUX-region evolution. Recent work has shown that autoimmunity is a major evolutionary constraint on anti-phage defence systems, with increased defence activity often accompanied by increased self-toxicity, thereby favouring regulatory mechanisms that limit basal activity (Aframian et al., 2025). We therefore speculate that the apparent coupling of DUF4062 catalytic activity to AUX-dependent activation provides an intrinsic layer of self-regulation that minimises basal toxicity and improves compatibility following horizontal transfer into diverse host backgrounds. Such a mechanism could make single-gene systems such as Evangelion less prone to autoimmunity than constitutively active or more complex multi-component systems, while preserving the capacity to evolve highly specialised sensing functions. This evolutionary logic could help explain both the widespread distribution of Evangelion systems and the pronounced lineage-specific differences in phage susceptibility observed across the family. The observation that Eva01 targets staphylococcal *Silviaviruses* whereas Eva03 interferes with *E. coli Dhillonviruses* further suggests that different Evangelion lineages have adapted to specific host-phage environments while retaining a common DUF4062-based framework.

Our findings also highlight the value of MGE perturbation as a complementary strategy for defence-system discovery. Most anti-phage systems have been discovered through guilt-by-association approaches that exploit co-localisation with known defence genes or defence islands (Doron et al., 2018; Bernheim and Sorek, 2020). By contrast, Eva01 was identified through a phenotype-first strategy in which removal of a naturally occurring plasmid altered phage susceptibility. This approach enabled the discovery of a single-gene defence system lacking obvious association with canonical defence loci and highlights how experimental interrogation of MGEs can uncover immune functions that remain invisible to current computational frameworks. Given the diversity of MGEs and their central role in shaping bacterial pan-immunity, systematic perturbation of plasmids, prophages and other mobile elements may provide a powerful route for identifying additional classes of antiviral systems that that remain inaccessible to current sequence-based discovery approaches.

In summary, Evangelion expands the known repertoire of mobile anti-phage systems and identifies antimicrobial-resistance plasmids as reservoirs of DUF4062-based immunity. Our work provides experimental validation of a previously uncharacterised DUF4062 defence family and establishes DUF4062 proteins as a new component of the bacterial pan-immune repertoire (Mordret et al., 2026; Osinski et al., 2026). Defining the biochemical activity of DUF4062 proteins, identifying the phage-derived signals that trigger Evangelion activation and understanding the evolutionary rules governing DUF4062-AUX compatibility represent important next steps towards understanding this defence family.

## Acknowledgements

Authors would like to thank Iñigo Lasa for *S. aureus* Mw2 pMw2 cured strain. We thank Mads F. Hansen, Andreas Haag, Malcolm White, and Janine Z. Bowring for helpful discussions and Andy Millard for helping with Nanopore sequencing of φMR003 escaper phages. We also thank Thorsten Gravert from Cmbio for his help with analysis and interpretation of the metabolomics data. Finally, we would like to thank Anette Hørdum Løth and Ayoe Lüchau for all their assistance in the laboratory.

This work was supported by research grants Villum Experiment (VIL58733-Weaponizable satellites) from Villum Foundation, and a JSPS Fellowship (PE24040) to R.I-C., JSPS KAKENHI (No. 25K21732) from Japan Society for the Promotion of Science (JSPS) to K.K.

## Author contributions

R.I-C. wrote the manuscript with input of all authors.

## Data and materials availability

All materials developed in this study will be made available upon request.

## Supplementary Figure legends

Figure S1. pMw2 protects *S. aureus* against *Silviavirus* infection across a range of phage burdens. Related to Figure 1. (A) Growth curves (*n*=3) of *S. aureus* Mw2 carrying pMw2 or a plasmid-cured derivative following infection with φMR003 at different multiplicities of infection (MOI). (B) Growth curves (*n*=3) following infection with φ74 at different MOIs.

Figure S2. Definition of the minimal functional Eva01 locus and complementation of defence phenotype in Mw2. Related to Figure 1. Representative spot assays of additional Eva01 constructs containing different combinations of upstream and downstream regions surrounding the DUF4062-containing gene. Constructs retaining the DUF4062 core and associated regulatory sequences confer protection against φMR003, whereas constructs lacking essential regions fail to protect against phage infection.

Figure S3. Inducible expression and functional mapping of Eva01. Related to Figure 1. Schematic of the tetracycline-inducible expression system used to define the minimal Eva01 locus. Representative spot assays performed in the presence and absence of anhydrotetracycline (aTc) demonstrate defence activity of constructs pEva01v1-v4. Protection is retained in constructs containing the minimal Eva01 region, consistent with basal expression from local regulatory elements.

Figure S4. Comparative genomic context of Eva01-containing plasmids. Related to Figure 1. Phylogenetic relationships and gene neighbourhoods of representative Eva01-positive plasmids identified from public databases. Conserved Eva01 loci are positioned adjacent to diverse combinations of antimicrobial-resistance genes, mobility-associated genes, toxin-antitoxin modules and hypothetical proteins, illustrating the diversity of Eva01-containing plasmid backbones.

Figure S5. Conservation of DUF4062 motifs across Evangelion proteins. Related to Figure 3. Multiple sequence alignments of representative Evangelion proteins from diverse bacterial hosts. Conserved residues and secondary-structure elements are indicated. The aromatic core motif, acidic-SYT region and DUF4062-specific Q/W/E motif are strongly conserved across the family. Residues selected for mutagenesis are highlighted.

Figure S6. Conservation of Evangelion-associated DUF4062 proteins. Related to Figure 3. Expanded sequence alignments and phylogenetic analysis of representative Evangelion homologues. Conserved DUF4062 residues remain identifiable despite extensive divergence in surrounding regions. The accompanying phylogeny illustrates the evolutionary relationships among representative homologues used throughout this study.

Figure S7. Additional metabolomic and structural analyses of Eva01. Related to Figure 4. (A) Representative chromatograms showing metabolite profiles in strains expressing wild-type or mutant Eva01 variants before and after φMR003 infection. (B-C) Quantification of additional nucleotide-associated metabolites altered during Evangelion activation. These data support widespread remodelling of nucleotide metabolism during Eva01-mediated defence.

Figure S8. Predicted oligomeric assemblies and native mass spectrometry of Eva01. Related to Figure 4. (A-B) AlphaFold-Multimer models of Eva01 DUF4062-containing regions in the presence of NAD⁺ or ADP. Predicted oligomeric assemblies display stacked interfaces involving conserved DUF4062 regions. (C) Native mass spectrometry reveals multiple species corresponding to monomeric, dimeric and higher-order assemblies, supporting the ability of Eva01 to self-associate in solution.

Figure S9. Additional characterization of phage escapers and replication-associated loci. Related to Figure 5. (A-B) Representative spot assays showing infection of Eva01-expressing strains by independent φMR003 and φ74 escaper mutants. (C) Comparative genomic analysis of additional *Silviavirus* genomes demonstrating conservation of the replication-associated locus targeted by escaper mutations. (D) Representative plating assays illustrating differential susceptibility of wild-type and mutant phages.

Figure S10. Expanded evolutionary analysis of DUF4062-containing defence systems. Related to Figure 6. (A) Sequence-similarity network of representative DUF4062-containing proteins showing clustering of Evangelion lineages, Swarożyc and TEP1-like proteins. (B) Domain-architecture comparison and structural overlays of representative DUF4062 proteins. Conserved DUF4062 cores are retained despite extensive diversification of auxiliary domains and overall protein architecture.

Figure S11. Conserved-domain relationships between Eva01 and TEP1-like proteins. Related to Figure 6 and Figure 7. Conserved-domain searches identifying DUF4062 as the shared feature between Eva01 and bacterial/eukaryotic TEP1-like proteins. Additional domains present in TEP1-like proteins, including TROVE, NACHT and WD40-repeat regions, are indicated.

Figure S12. Structural conservation of DUF4062 regions from Evangelion and TEP1 proteins. Related to Figure 7. Multiple sequence alignment and secondary-structure comparison of DUF4062-containing regions from Evangelion proteins, TEP1-like proteins and representative homologues. Conserved residues associated with the predicted DUF4062 fold are highlighted, supporting an evolutionary relationship between bacterial Evangelion systems and eukaryotic TEP1-like proteins.

