## Supplementary Figure for "Antimicrobial-resistance plasmids encode Evangelion, a widespread DUF4062 anti-phage defence family" (from another version of this preprint: version 1, 2026-06-26)

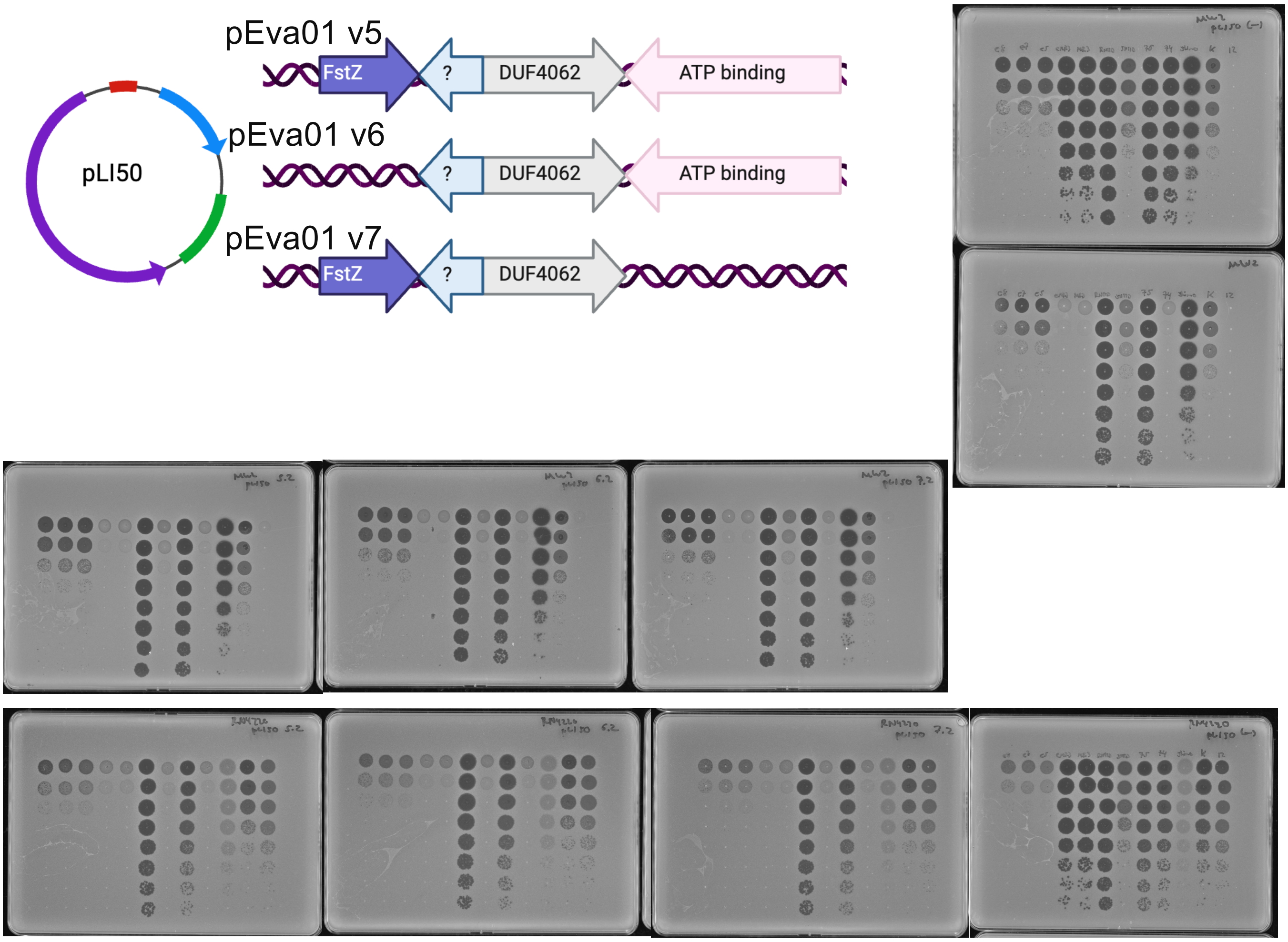
pEva01 v5
FstZ
?
DUF4062
ATP binding
pLI50
pEva01 v6
?
DUF4062
ATP binding
pEva01 v7
FstZ
?
DUF4062

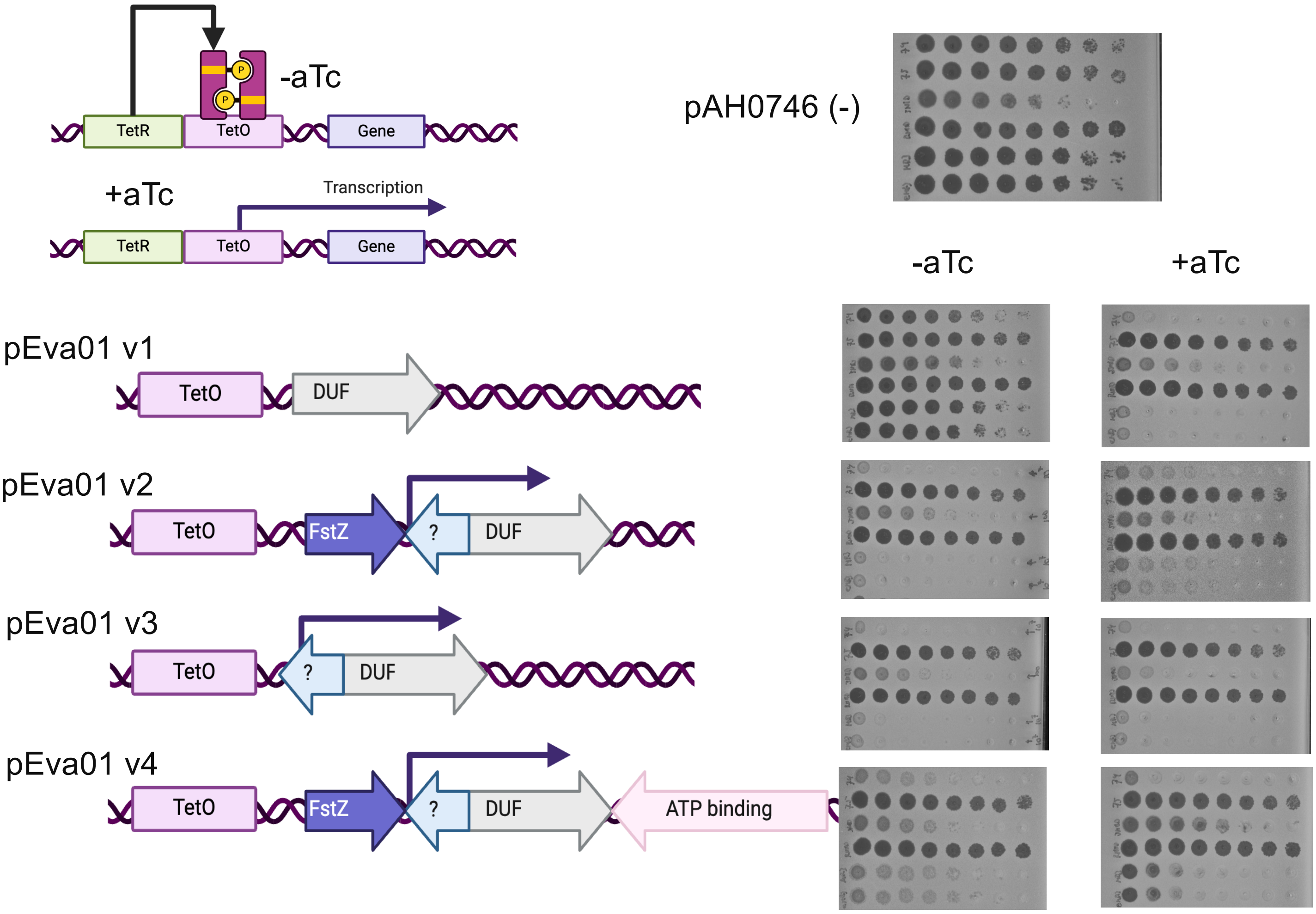

-aTc
TetR
TetO
Gene
+aTc
Transcription
TetR
TetO
Gene
pAH0746 (-)
-aTc
+aTc
pEva01 v1
TetO
DUF
pEva01 v2
TetO
FstZ
?
DUF
pEva01 v3
TetO
?
DUF
pEva01 v4
TetO
FstZ
?
DUF
ATP binding

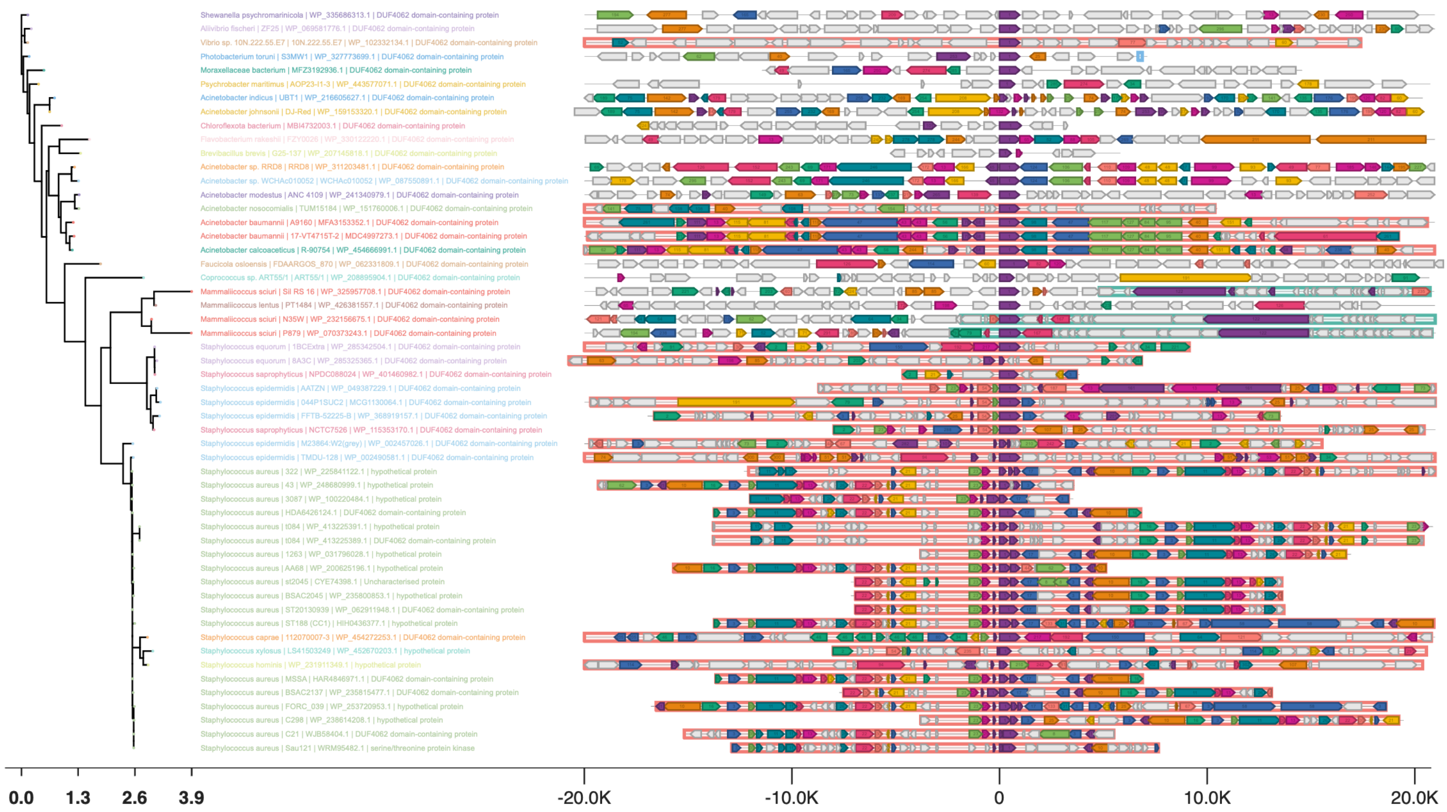

Shewanella psychromarinicola | WP_335686313.1 | DUF4062 domain-containing protein
Aliivibrio fischeri | ZF25 | WP_069581776.1 | DUF4062 domain-containing protein
Vibrio sp. 10N.222.55.E7 | 10N.222.55.E7 | WP_102332134.1 | DUF4062 domain-containing protein
Photobacterium toruni | S3MW1 | WP_327773699.1 | DUF4062 domain-containing protein
Moraxellaceae bacterium | MFZ3192936.1 | DUF4062 domain-containing protein
Psychrobacter maritimus | AOP23-I1-3 | WP_443577071.1 | DUF4062 domain-containing protein
Acinetobacter indicus | UBT1 | WP_216605627.1 | DUF4062 domain-containing protein
Acinetobacter johnsonii | DJ-Red | WP_159153320.1 | DUF4062 domain-containing protein
Chloroflexota bacterium | MBI4732003.1 | DUF4062 domain-containing protein
Flavobacterium rakeshii | FZY0026 | WP_330122220.1 | DUF4062 domain-containing protein
Brevibacillus brevis | G25-137 | WP_207145818.1 | DUF4062 domain-containing protein
Acinetobacter sp. RRD8 | RRD8 | WP_311203481.1 | DUF4062 domain-containing protein
Acinetobacter sp. WCHAc010052 | WCHAc010052 | WP_087550891.1 | DUF4062 domain-containing protein
Acinetobacter modestus | ANC 4109 | WP_241340979.1 | DUF4062 domain-containing protein
Acinetobacter nosocomialis | TUM15184 | WP_151760006.1 | DUF4062 domain-containing protein
Acinetobacter baumannii | A9160 | MFA3153352.1 | DUF4062 domain-containing protein
Acinetobacter baumannii | 17-VT4715T-2 | MDC4997273.1 | DUF4062 domain-containing protein
Acinetobacter calcoaceticus | R-90754 | WP_454666991.1 | DUF4062 domain-containing protein
Faucicola osloensis | FDAARGOS_870 | WP_062331809.1 | DUF4062 domain-containing protein
Coprococcus sp. ART55/1 | ART55/1 | WP_208895904.1 | DUF4062 domain-containing protein
Mammaliicoccus sciuri | Sil RS 16 | WP_325957708.1 | DUF4062 domain-containing protein
Mammaliicoccus lentus | PT1484 | WP_426381557.1 | DUF4062 domain-containing protein
Mammaliicoccus sciuri | N35W | WP_232156675.1 | DUF4062 domain-containing protein
Mammaliicoccus sciuri | P879 | WP_070373243.1 | DUF4062 domain-containing protein
Staphylococcus equorum | 1BCExtra | WP_285342504.1 | DUF4062 domain-containing protein
Staphylococcus equorum | 8A3C | WP_285325365.1 | DUF4062 domain-containing protein
Staphylococcus saprophyticus | NPDC088024 | WP_401460982.1 | DUF4062 domain-containing protein
Staphylococcus epidermidis | AATZN | WP_049387229.1 | DUF4062 domain-containing protein
Staphylococcus epidermidis | 044P1SUC2 | MCG1130064.1 | DUF4062 domain-containing protein
Staphylococcus epidermidis | FFTB-52225-B | WP_368919157.1 | DUF4062 domain-containing protein
Staphylococcus saprophyticus | NCTC7526 | WP_115353170.1 | DUF4062 domain-containing protein
Staphylococcus epidermidis | M23864:W2(grey) | WP_002457026.1 | DUF4062 domain-containing protein
Staphylococcus epidermidis | TMDU-128 | WP_002490581.1 | DUF4062 domain-containing protein
Staphylococcus aureus | 322 | WP_225841122.1 | hypothetical protein
Staphylococcus aureus | 43 | WP_248680999.1 | hypothetical protein
Staphylococcus aureus | 3087 | WP_100220484.1 | hypothetical protein
Staphylococcus aureus | HDA6426124.1 | DUF4062 domain-containing protein
Staphylococcus aureus | t084 | WP_413225391.1 | hypothetical protein
Staphylococcus aureus | t084 | WP_413225389.1 | DUF4062 domain-containing protein
Staphylococcus aureus | 1263 | WP_031796028.1 | hypothetical protein
Staphylococcus aureus | AA68 | WP_200625196.1 | hypothetical protein
Staphylococcus aureus | st2045 | CYE74398.1 | Uncharacterised protein
Staphylococcus aureus | BSAC2045 | WP_235800853.1 | hypothetical protein
Staphylococcus aureus | ST20130939 | WP_062911948.1 | DUF4062 domain-containing protein
Staphylococcus aureus | ST188 (CC1) | HIH0436377.1 | hypothetical protein
Staphylococcus caprae | 112070007-3 | WP_454272253.1 | DUF4062 domain-containing protein
Staphylococcus xylosus | LS41503249 | WP_452670203.1 | hypothetical protein
Staphylococcus hominis | WP_231911349.1 | hypothetical protein
Staphylococcus aureus | MSSA | HAR4846971.1 | DUF4062 domain-containing protein
Staphylococcus aureus | BSAC2137 | WP_235815477.1 | DUF4062 domain-containing protein
Staphylococcus aureus | FORC_039 | WP_253720953.1 | hypothetical protein
Staphylococcus aureus | C298 | WP_238614208.1 | hypothetical protein
Staphylococcus aureus | C21 | WJB58404.1 | DUF4062 domain-containing protein
Staphylococcus aureus | Sau121 | WRM95482.1 | serine/threonine protein kinase
0.0
1.3
2.6
3.9
-20.0K
-10.0K
0
10.0K
20.0K

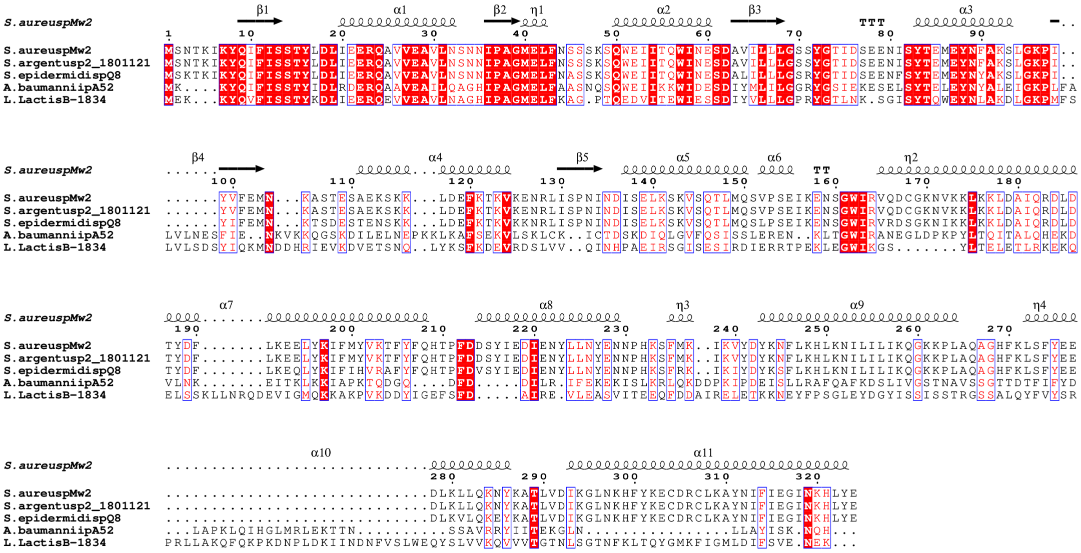
S.aureuspMw2
β1 α1 β2 η1 α2 β3 TTT α3
S.aureuspMw2
S.argentusp2_1801121
S.epidermidispQ8
A.baumanniipA52
L.LactisB-1834
β4 α4 β5 α5 α6 TT η2
α7 α8 η3 α9 η4
α10 α11

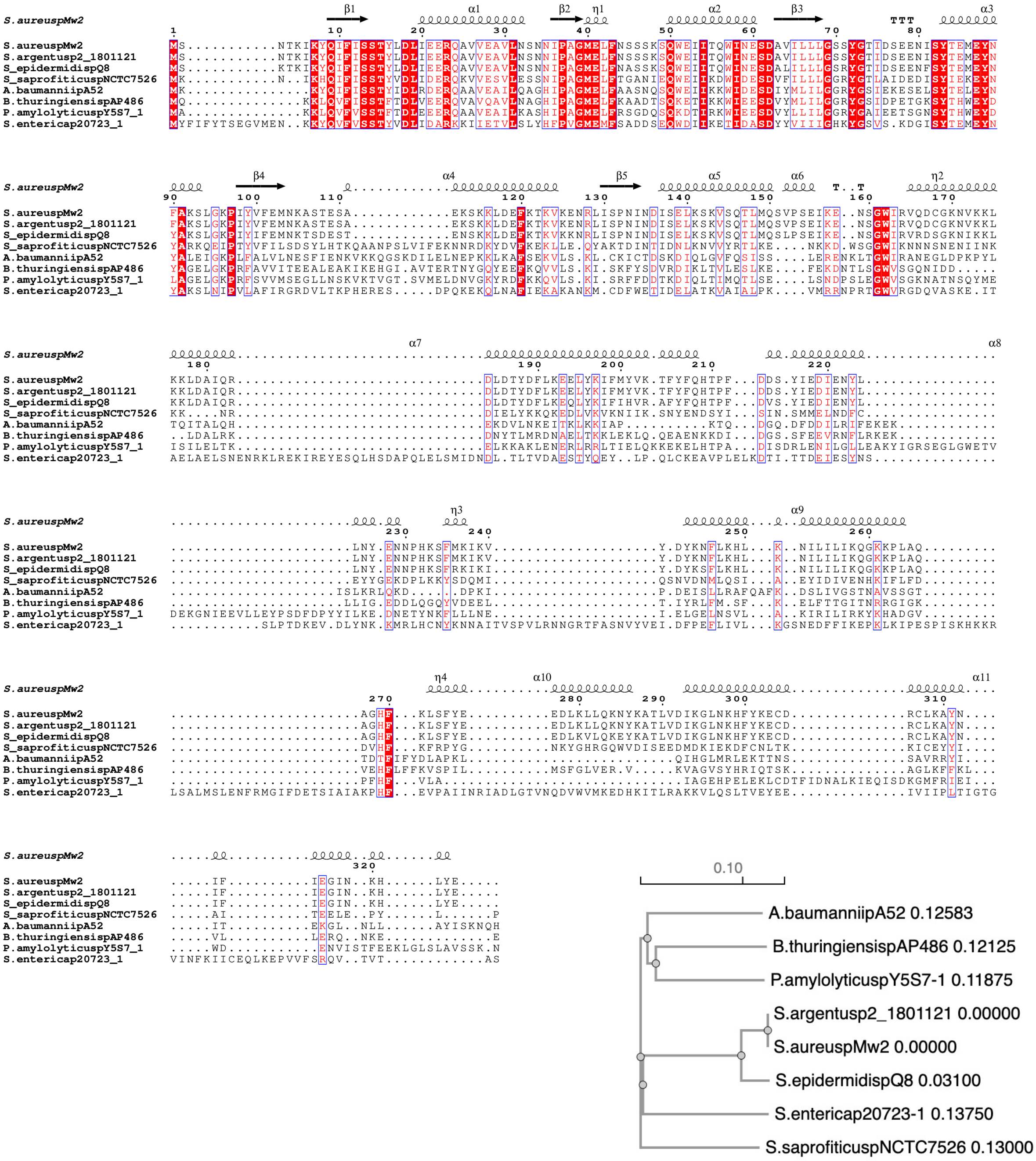
S.aureuspMw2
S.argentusp2_1801121
S_epidermidispQ8
S_saprofiticuspNCTC7526
A.baumanniipA52
B.thuringiensispAP486
P.amylolyticuspY5S7_1
S.entericap20723_1
0.10
A.baumanniipA52 0.12583
B.thuringiensispAP486 0.12125
P.amylolyticuspY5S7-1 0.11875
S.argentusp2_1801121 0.00000
S.aureuspMw2 0.00000
S.epidermidispQ8 0.03100
S.entericap20723-1 0.13750
S.saprofiticuspNCTC7526 0.13000

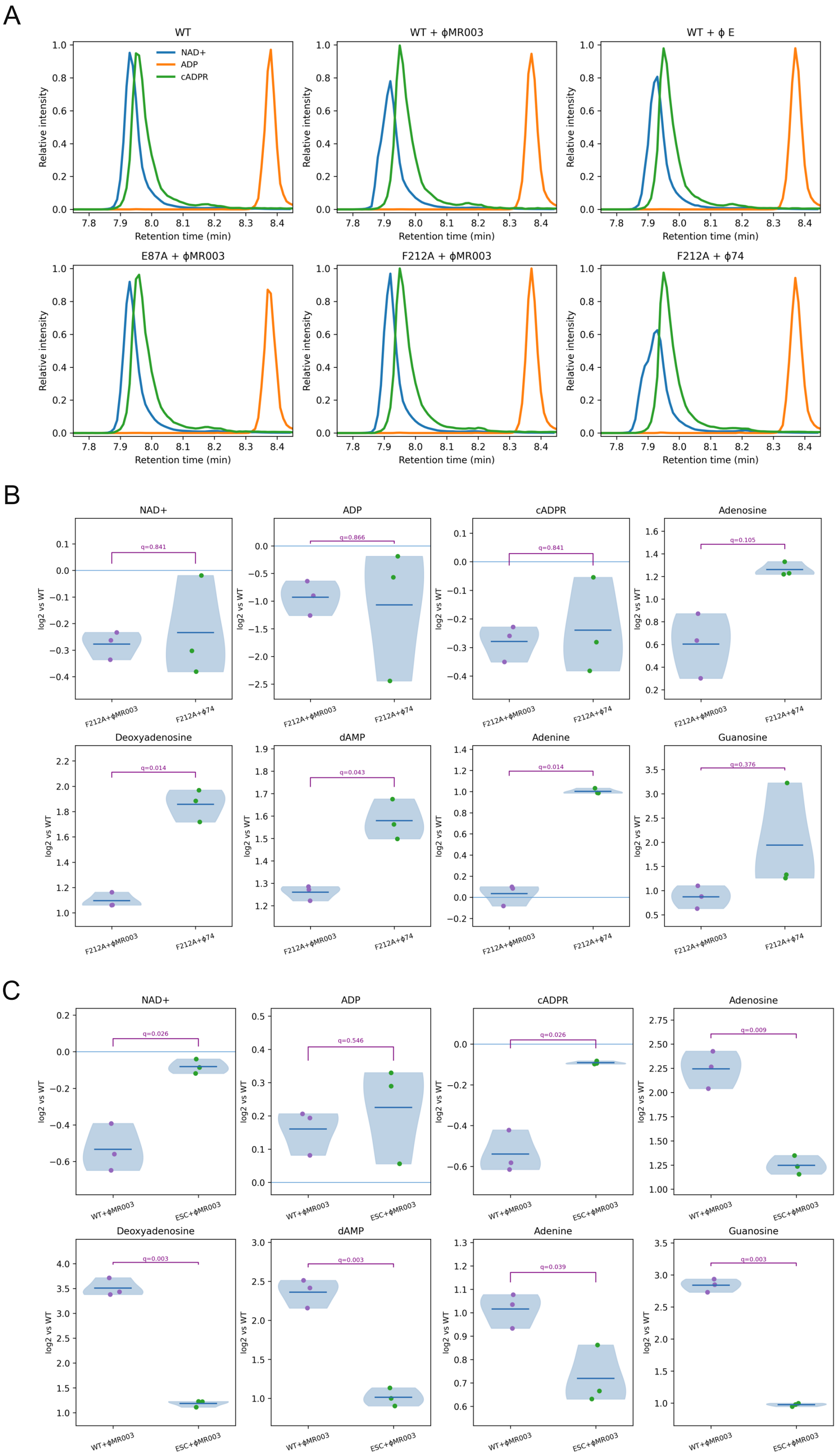
A
WT
WT + ϕMR003
WT + ϕ E
E87A + ϕMR003
F212A + ϕMR003
F212A + ϕ74
NAD+
ADP
cADPR
Relative intensity
Retention time (min)
B
NAD+
ADP
cADPR
Adenosine
Deoxyadenosine
dAMP
Adenine
Guanosine
q=0.841
q=0.866
q=0.841
q=0.105
q=0.014
q=0.043
q=0.014
q=0.376
log2 vs WT
F212A+ϕMR003
F212A+ϕ74
C
NAD+
ADP
cADPR
Adenosine
Deoxyadenosine
dAMP
Adenine
Guanosine
q=0.026
q=0.546
q=0.026
q=0.009
q=0.003
q=0.003
q=0.039
q=0.003
log2 vs WT
WT+ϕMR003
ESC+ϕMR003

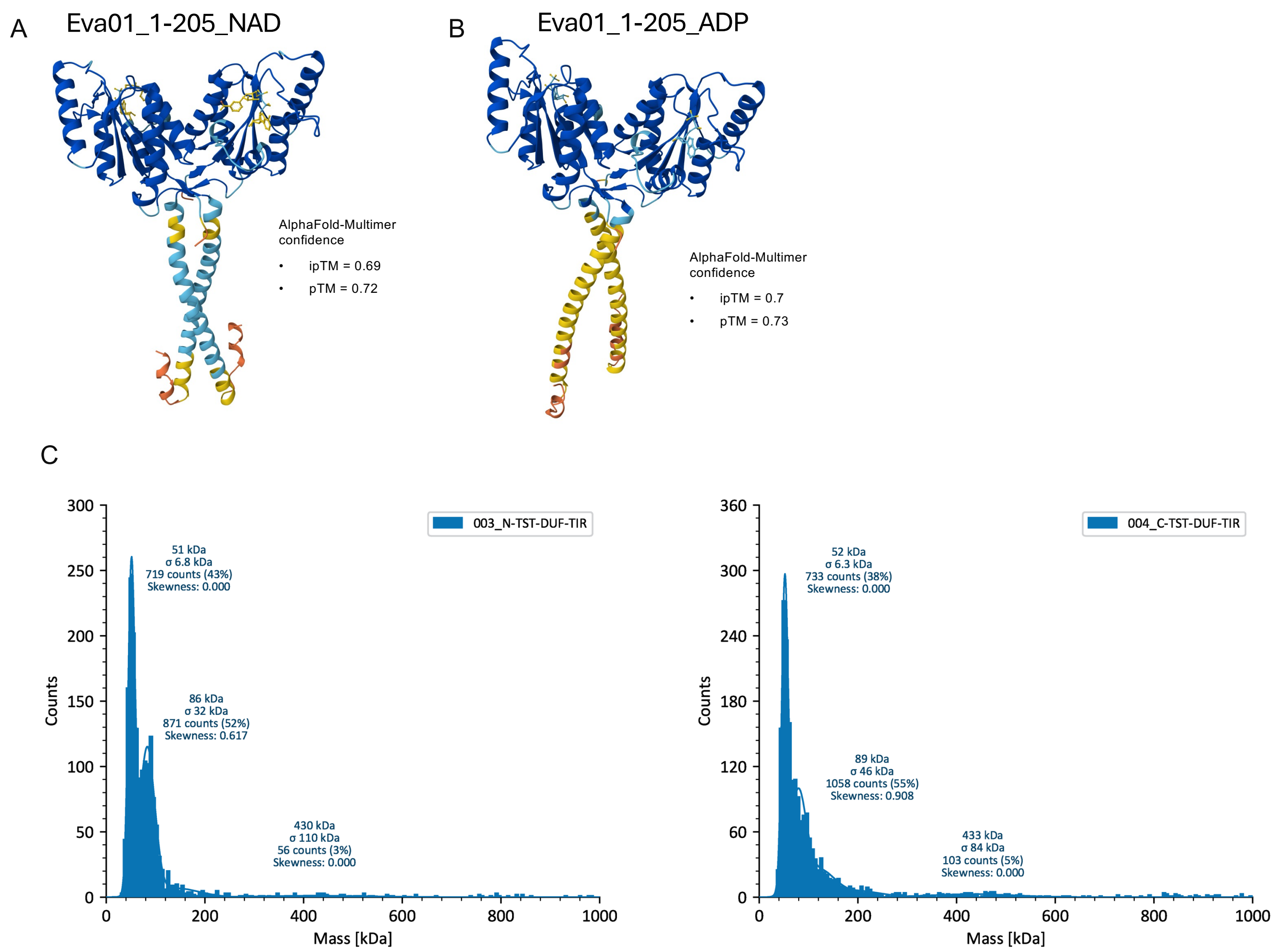
A
Eva01_1-205_NAD
AlphaFold-Multimer confidence
• ipTM = 0.69
• pTM = 0.72
B
Eva01_1-205_ADP
AlphaFold-Multimer confidence
• ipTM = 0.7
• pTM = 0.73
C
003_N-TST-DUF-TIR
51 kDa
σ 6.8 kDa
719 counts (43%)
Skewness: 0.000
86 kDa
σ 32 kDa
871 counts (52%)
Skewness: 0.617
430 kDa
σ 110 kDa
56 counts (3%)
Skewness: 0.000
Counts
Mass [kDa]
004_C-TST-DUF-TIR
52 kDa
σ 6.3 kDa
733 counts (38%)
Skewness: 0.000
89 kDa
σ 46 kDa
1058 counts (55%)
Skewness: 0.908
433 kDa
σ 84 kDa
103 counts (5%)
Skewness: 0.000
Counts
Mass [kDa]

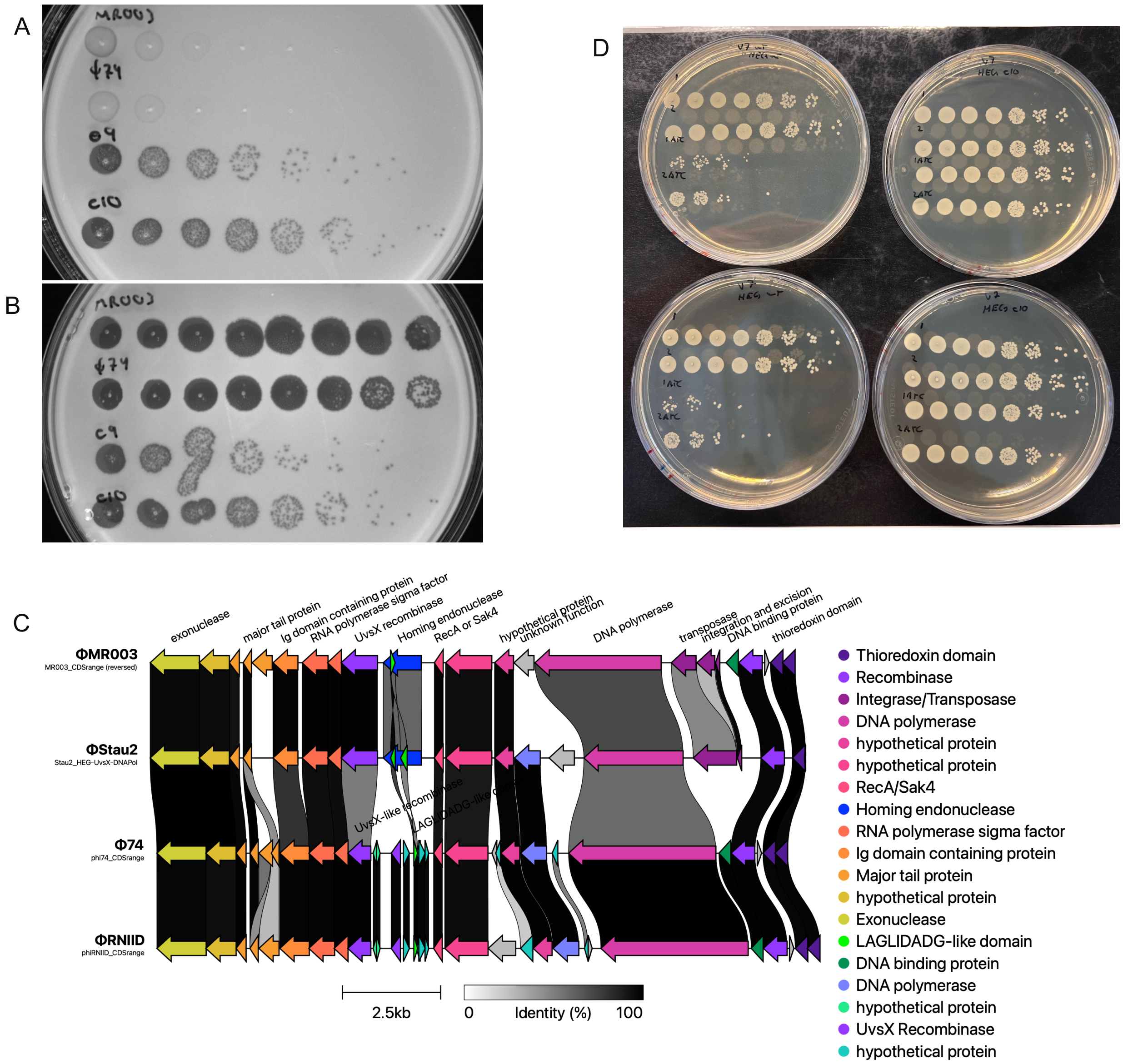
A
MR003
Φ74
c9
c10
B
MR003
Φ74
c9
c10
C
ΦMR003
MR003_CDSrange (reversed)
ΦStau2
Stau2_HEG-UvsX-DNAPol
Φ74
phi74_CDSrange
ΦRNIID
phiRNIID_CDSrange
exonuclease
major tail protein
Ig domain containing protein
RNA polymerase sigma factor
UvsX recombinase
Homing endonuclease
RecA or Sak4
hypothetical protein
unknown function
DNA polymerase
transposase
integration and excision
DNA binding protein
thioredoxin domain
UvsX-like recombinase
LAGLIDADG-like
2.5kb
0
Identity (%)
100
Thioredoxin domain
Recombinase
Integrase/Transposase
DNA polymerase
hypothetical protein
hypothetical protein
RecA/Sak4
Homing endonuclease
RNA polymerase sigma factor
Ig domain containing protein
Major tail protein
hypothetical protein
Exonuclease
LAGLIDADG-like domain
DNA binding protein
DNA polymerase
hypothetical protein
UvsX Recombinase
hypothetical protein
D

A

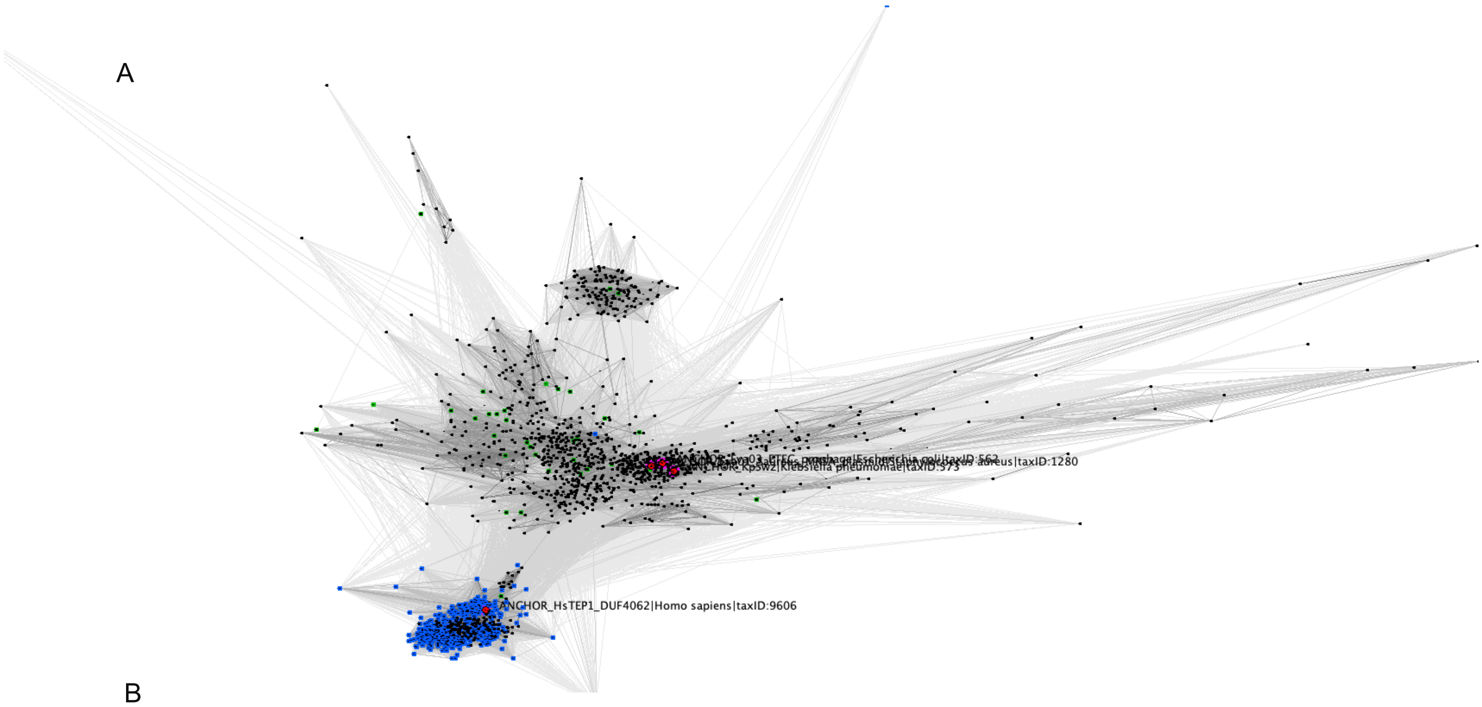
ANCHOR_HsTEP1_DUF4062|Homo sapiens|taxID:9606

B

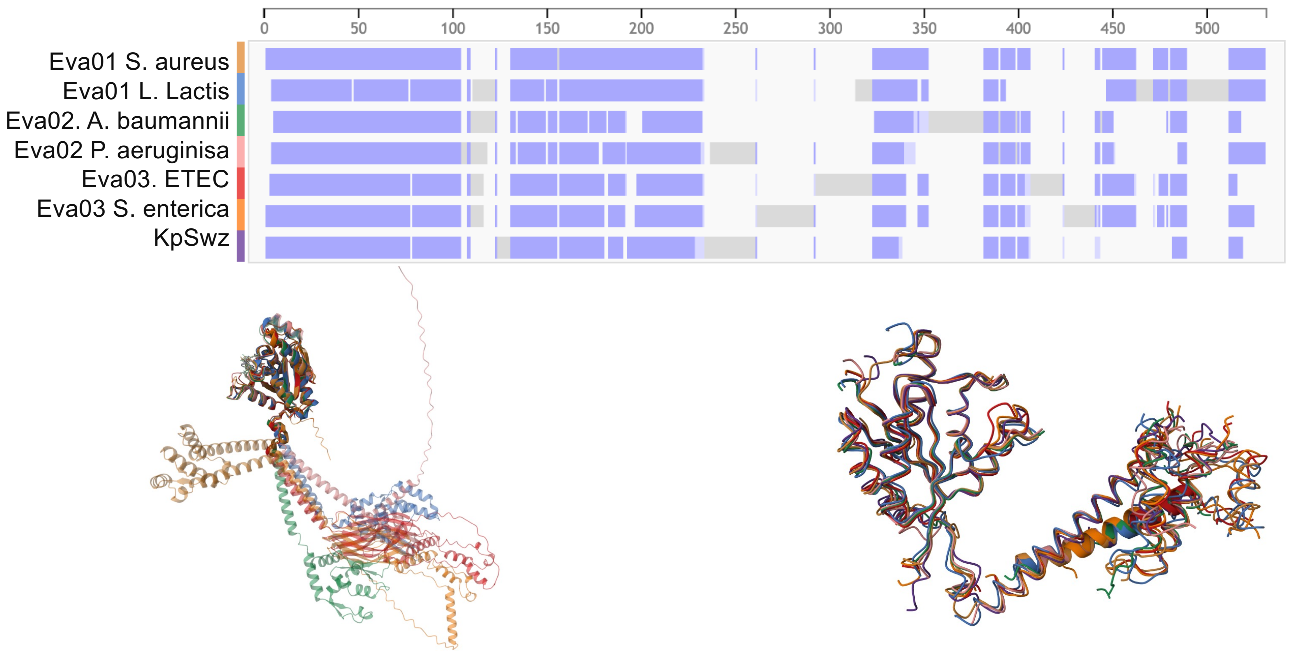
0
50
100
150
200
250
300
350
400
450
500
Eva01 S. aureus
Eva01 L. Lactis
Eva02. A. baumannii
Eva02 P. aeruginisa
Eva03. ETEC
Eva03 S. enterica
KpSwz

A

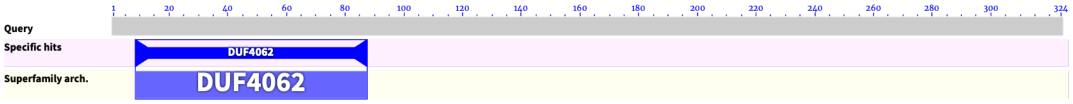

Query
Specific hits
Superfamily arch.
DUF4062
DUF4062


B

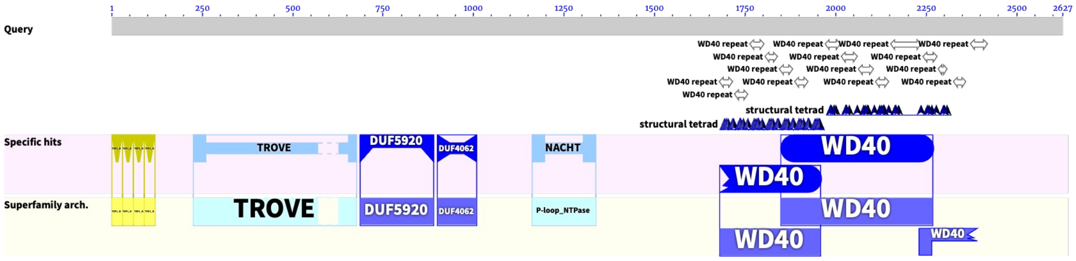

Query
Specific hits
Superfamily arch.
WD40 repeat
structural tetrad
TROVE
DUF5920
DUF4062
NACHT
WD40
P-loop_NTPase

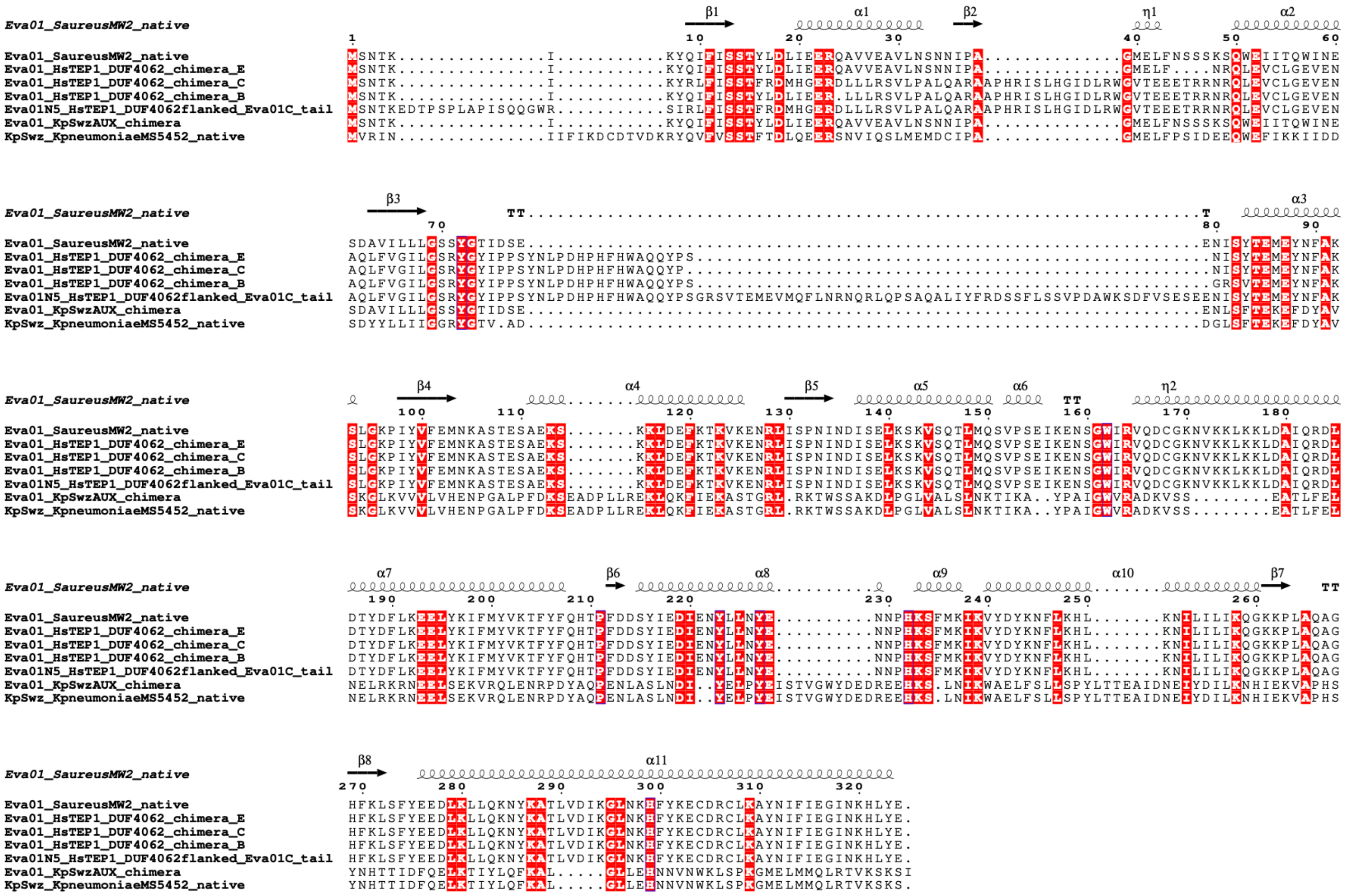

Eva01_SaureusMW2_native
β1 α1 β2 η1 α2
1 10 20 30 40 50 60
Eva01_SaureusMW2_native MSNTK...............I...........KYQIFISSTYLDLIEERQAVVEAVLNSNNIPA..............GMELFNSSSKSQWEIITQWINE
Eva01_HsTEP1_DUF4062_chimera_E MSNTK...............I...........KYQIFISSTYLDLIEERQAVVEAVLNSNNIPA..............GMELF....NRQLEVCLGEVEN
Eva01_HsTEP1_DUF4062_chimera_C MSNTK...............I...........KYRLFISSTFRDMHGERDLLLRSVLPALQARAAPHRISLHGIDLRWGVTEETRRNRQLEVCLGEVEN
Eva01_HsTEP1_DUF4062_chimera_B MSNTK...............I...........KYQIFISSTYLDLIEER.LLLRSVLPALQARAAPHRISLHGIDLRWGVTEETRRNRQLEVCLGEVEN
Eva01N5_HsTEP1_DUF4062flanked_Eva01C_tail MSNTKEDTPSPLAPISQQGWR...........SIRLFISSTFRDMHGERDLLLRSVLPALQARAAPHRISLHGIDLRWGVTEETRRNRQLEVCLGEVEN
Eva01_KpSwzAUX_chimera MSNTK...............I...........KYQIFISSTYLDLIEERQAVVEAVLNSNNIPA..............GMELFNSSSKSQWEIITQWINE
KpSwz_KpneumoniaeMS5452_native MVRIN...............IIFIKDCDTVDKRYQVFVSSTFTDLQEERSNVIQSLMEMDCIPA..............GMELFPSIDEEQWEFIKKIIDD
Eva01_SaureusMW2_native
β3 TT T α3
70 80 90
Eva01_SaureusMW2_native SDAVILLLGSSYGTIDSE...............................................................ENISYTEMEYNFAK
Eva01_HsTEP1_DUF4062_chimera_E AQLFVGILGSRYGYIPPSYNLPDHPHFHWAQQYPS..............................................NISYTEMEYNFAK
Eva01_HsTEP1_DUF4062_chimera_C AQLFVGILGSRYGYIPPSYNLPDHPHFHWAQQYP...............................................NISYTEMEYNFAK
Eva01_HsTEP1_DUF4062_chimera_B AQLFVGILGSRYGYIPPSYNLPDHPHFHWAQQYPS..............................................GRSVTEMEYNFAK
Eva01N5_HsTEP1_DUF4062flanked_Eva01C_tail AQLFVGILGSRYGYIPPSYNLPDHPHFHWAQQYPSGRSVTEMEVMQFLNRNQRLQPSAQALIYFRDSSFLSSVPDAWKSDFVSESEENISYTEMEYNFAK
Eva01_KpSwzAUX_chimera SDAVILLLGSSYGTIDSE...............................................................ENLSFTEKEFDYAV
KpSwz_KpneumoniaeMS5452_native SDYYLLIIGGRYGTV.AD...............................................................DGLSFTEKEFDYAV
Eva01_SaureusMW2_native
β4 α4 β5 α5 α6 TT η2
100 110 120 130 140 150 160 170 180
Eva01_SaureusMW2_native SLGKPIYVFEMNKASTESAEKS.......KKIDEFKTKVKENRLISPNINDISELKSKVSQTLMQSVPSEIKENSGWIRVQDCGKNVKKLKKLDAIQRDL
Eva01_HsTEP1_DUF4062_chimera_E SLGKPIYVFEMNKASTESAEKS.......KKIDEFKTKVKENRLISPNINDISELKSKVSQTLMQSVPSEIKENSGWIRVQDCGKNVKKLKKLDAIQRDL
Eva01_HsTEP1_DUF4062_chimera_C SLGKPIYVFEMNKASTESAEKS.......KKIDEFKTKVKENRLISPNINDISELKSKVSQTLMQSVPSEIKENSGWIRVQDCGKNVKKLKKLDAIQRDL
Eva01_HsTEP1_DUF4062_chimera_B SLGKPIYVFEMNKASTESAEKS.......KKIDEFKTKVKENRLISPNINDISELKSKVSQTLMQSVPSEIKENSGWIRVQDCGKNVKKLKKLDAIQRDL
Eva01N5_HsTEP1_DUF4062flanked_Eva01C_tail SLGKPIYVFEMNKASTESAEKS.......KKIDEFKTKVKENRLISPNINDISELKSKVSQTLMQSVPSEIKENSGWIRVQDCGKNVKKLKKLDAIQRDL
Eva01_KpSwzAUX_chimera SKGLKVVVLVHENPGALPFDKSEADPLLREKLQKFIEKASTGRL.RKTWSSAKDLPGLVALSLNKTIKA..YPAIGWVRADKVSS........EATLFEL
KpSwz_KpneumoniaeMS5452_native SKGLKVVVLVHENPGALPFDKSEADPLLREKLQKFIEKASTGRL.RKTWSSAKDLPGLVALSLNKTIKA..YPAIGWVRADKVSS........EATLFEL
Eva01_SaureusMW2_native
α7 β6 α8 α9 α10 β7 TT
190 200 210 220 230 240 250 260
Eva01_SaureusMW2_native DTYDFLKEELYKIFMYVKTFYFQHTPFDDSYIEDIENYLLNYE..........NNPHKSFMKIKVYDYKNFLKHL.......KNILILIKQGKKPLAQAG
Eva01_HsTEP1_DUF4062_chimera_E DTYDFLKEELYKIFMYVKTFYFQHTPFDDSYIEDIENYLLNYE..........NNPHKSFMKIKVYDYKNFLKHL.......KNILILIKQGKKPLAQAG
Eva01_HsTEP1_DUF4062_chimera_C DTYDFLKEELYKIFMYVKTFYFQHTPFDDSYIEDIENYLLNYE..........NNPHKSFMKIKVYDYKNFLKHL.......KNILILIKQGKKPLAQAG
Eva01_HsTEP1_DUF4062_chimera_B DTYDFLKEELYKIFMYVKTFYFQHTPFDDSYIEDIENYLLNYE..........NNPHKSFMKIKVYDYKNFLKHL.......KNILILIKQGKKPLAQAG
Eva01N5_HsTEP1_DUF4062flanked_Eva01C_tail DTYDFLKEELYKIFMYVKTFYFQHTPFDDSYIEDIENYLLNYE..........NNPHKSFMKIKVYDYKNFLKHL.......KNILILIKQGKKPLAQAG
Eva01_KpSwzAUX_chimera NELRKRNEELSEKVRQLENRPDYAQPENLASLNDI..YELPYEISTVGWYDEDREEHKS.LNIKWAELFSLLSPYLTTEAIDNEIYDILKNHIEKVAPHS
KpSwz_KpneumoniaeMS5452_native NELRKRNEELSEKVRQLENRPDYAQPENLASLNDI..YELPYEISTVGWYDEDREEHKS.LNIKWAELFSLLSPYLTTEAIDNEIYDILKNHIEKVAPHS
Eva01_SaureusMW2_native
β8 α11
270 280 290 300 310 320
Eva01_SaureusMW2_native HFKLSFYEEDLKLLQKNYKATLVDIKGLNKHFYKECDRCLKAYNIFIEGINKHLYE.
Eva01_HsTEP1_DUF4062_chimera_E HFKLSFYEEDLKLLQKNYKATLVDIKGLNKHFYKECDRCLKAYNIFIEGINKHLYE.
Eva01_HsTEP1_DUF4062_chimera_C HFKLSFYEEDLKLLQKNYKATLVDIKGLNKHFYKECDRCLKAYNIFIEGINKHLYE.
Eva01_HsTEP1_DUF4062_chimera_B HFKLSFYEEDLKLLQKNYKATLVDIKGLNKHFYKECDRCLKAYNIFIEGINKHLYE.
Eva01N5_HsTEP1_DUF4062flanked_Eva01C_tail HFKLSFYEEDLKLLQKNYKATLVDIKGLNKHFYKECDRCLKAYNIFIEGINKHLYE.
Eva01_KpSwzAUX_chimera YNHTTIDFQELKTIYLQFKAL.....GLLEHNNVNWKLSPKGMELMMQLRTVKSKSI
KpSwz_KpneumoniaeMS5452_native YNHTTIDFQELKTIYLQFKAL.....GLLEHNNVNWKLSPKGMELMMQLRTVKSKS.
